# Feature-specific environmental contributions to human cortical architecture revealed by twin brainprints

**DOI:** 10.1101/2025.03.06.641380

**Authors:** Kristine B Walhovd, Anne Cecilie Sjøli Bråthen, Knut Overbye, Jonas Kransberg, Øystein Sørensen, Pablo F. Garrido, Inge K. Amlien, Jose-Luis Alatorre-Warren, Athanasia M. Mowinckel, Maksim Slivka, Nikolai O. Czajkowski, Yunpeng Wang, Paulina Due-Tønnessen, Jennifer R. Harris, Martin Lövdén, Didac Vidal-Pineiro, Lars Nyberg, Anders M. Fjell, Markus H. Sneve

## Abstract

How environmental variation shapes the human cerebral cortex remains incompletely understood. We compared cortical *brainprints* based in 210 twins (71 MZ and 34 DZ pairs, age 16-78 years) to distinguish prenatal, adult naturalistic and experimentally induced environmental variation from genetic contributions. Genetic effects were reflected in higher *brainprint* similarity within monozygotic (MZ) than dizygotic (DZ) pairs. Early environmental contributions were evident in lower brainprint similarity in MZ twin pairs with larger birthweight discordances, driven by area across the cortical ribbon. Later within-pair environmental differences in adult weight and lifestyle had minimal influence. Still, a 10-week virtual-reality navigation intervention revealed training-induced changes in the gray-white interface, with curvature and area changes supported by microstructural reconfigurations. In support of gene-environment interactions, relative brainprint similarity increased in MZ but diverged in DZ pairs following training. The results demonstrate that in adulthood, early life environmental difference persistently contributes to make the cortical architecture of genetically identical twins deviate. Environmental influence in adulthood in the form of training can still affect similarity of twin brainprints at the grey-white-matter boundary. These findings show that distinct environmental influences at prenatal and adult stage are differentially expressed across cortical features within a genetically informed framework.

## Introduction

Mammals have a genetic program for corticogenesis, much of which unfolds in utero ^1, 2^. Cortical architecture is thus highly heritable and vulnerable to prenatal genetic and environmental perturbations^1, 3–6^. Still, prominent cortical changes are normally seen throughout the human lifespan, and individual differences in the human brain can also be shaped by numerous experiences in life^7–14^. This scenario makes it a paramount challenge to determine the nature of influences on the developing, adult and aging human brain.

Multiple diseases affecting mental function, both neurodevelopmental and neurodegenerative, are associated with smaller cortical volumes^15–17^. Such individual differences are often ascribed to factors influencing trajectories of cortical change, and there is optimism for the possibility of improving brain health by environmental modification even in older adulthood ^18, 19^. However, neurodevelopmental origins of variation in the brain and its function through the lifespan are acknowledged^20–23^, and genetic factors in brain changes have been identified^24^. These perspectives on human brain dynamics need integration^25^. How and when can environmental influences, beyond genetics, change the human cerebral cortex? Comparisons of monozygotic (MZ) and dizygotic (DZ) twins allow separation of environmental and genetic influences, and within-pair comparison of MZ twins can help elucidate the impact of critical environmental effects on the brain while controlling for genetic background effects^26, 27^. Here we seek to identify which features of the adult and aging cortex may be ascribed to early versus later environmental influences, by longitudinally studying systematic variation in prenatal versus adult environment in genetically identical and fraternal twins in a combined observational-experimental design.

We examine how human cortical architecture – surface area, thickness, volume, and curvature holistically and feature-wise are related to different environmental factors and interrogate effects further using measures of cortical microstructure. Past theoretical and empirical work suggests that the impact of environmental events varies depending on the nature and timing of influences and among distinct cortical features^2, 4, 8–10, 21–23, 25, 28–34^. For instance, early environmental influences are likely to have a prominent influence on cortical surface area^2, 21–23, 35, 36^, whereas other cortical features may be more sensitive to later influences^9–14, 37^.

A *brainprint* ^38–40^ (Fig. 1) was computed from 272 cortical features (68 each for area, thickness, volume, and curvature^41, 42^) and used as an index of structural brain similarity of MZ and DZ twins. Brainprint similarity was expected to scale with genetic relatedness (i.e. be higher in MZ than in DZ) but be attenuated by variation in early environmental events reflected by birth weight differences. Thus, throughout adulthood, brainprints were predicted to be more similar for MZ twin pairs with minor birth weight discordances, and less similar within MZ twin pairs with more marked birth weight discordances.

**Figure 1.**
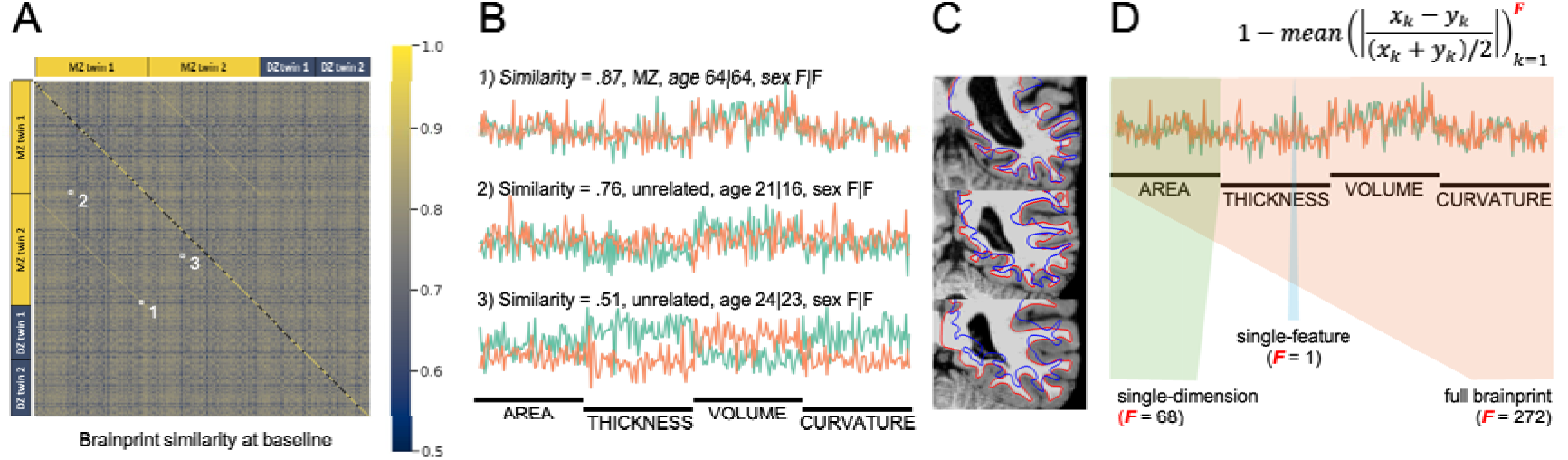
Anatomical brainprints and their similarity across individuals. A) Pairwise similarity between full brainprints for all 210 participants. Brainprints comprise 272 cortical features (area, thickness, volume, and curvature across 68 regions), here residualized for age (smooth) and sex to remove broad demographic effects. The main diagonal shows self-similarity for participants with two non-training sessions separated by 10 weeks and is marked in black where such data are unavailable. Within this subset, cross-session identification of individuals was perfect (127/127 correct; permutation p = 0.0001). Off-diagonal blocks of elevated similarity mark MZ twin pairs, whereas DZ similarity is weaker and not apparent as similarly distinct blocks. Brainprint similarity enabled identification of MZ co-twins (top-1 accuracy = 0.84; top-2 = 0.89; chance = 1/209; permutation p = 0.0001), while DZ similarity was statistically detectable but substantially weaker (top-1|top-2 = 0.13|0.21; permutation p = 0.0001). B) Example brainprints. Top: a highly similar MZ twin pair; middle: an unrelated pair with median similarity; bottom: the least similar pair in the sample. Each line is one individual’s normalized feature vector (272 features). These examples illustrate the practical similarity range, from near-identical (∼1 for repeated scans of the same individual) to typical unrelated values (∼0.5–0.75; see Figure S1 for full distributions). C) Corresponding cortical anatomy for the same pairs, shown by overlaying white and pial surfaces after rigid alignment to a common reference. These images serve only as intuition for anatomical correspondence; surface overlays are not used to compute brainprints. D) Schematic of the similarity metric. For each feature k, values x_k_ and y_k_ from two individuals are compared via their relative difference normalized by their average; similarity equals one minus the mean of these differences across all F features. This formulation handles heterogeneous feature scales and can be extended to reduced brainprints, including single features.

Adult environmental similarity was proxied by quantifying variation in factors such as BMI, blood pressure, alcohol and smoking, sleep^43^, physical activity^44^, and educational attainment. Given identical genes, differences in such variables reflect purely environmental factors in MZ twins. Based on the fetal origin of cortical architecture, variation in naturally occurring adult experiences was predicted to influence cortical characteristics to a lesser degree than early events.

Furthermore, the impact of a controlled environmental intervention in adulthood on brainprint similarity in MZ and DZ twins was examined using an experimental longitudinal virtual reality navigation training paradigm. Navigation training was chosen as a suitable paradigm for studying effects of controlled environment in interaction with genetic factors, as navigation ability shows wide variability related to both genetic and environmental differences^45, 46^, and effects of navigation training on brain structural characteristics have previously been demonstrated^11, 47^. An immersive virtual reality setting with real self-induced motion during navigation by bike was deemed appropriate to increase ecological validity, and was further motivated by previously demonstrated cortical volume and thickness effects in adulthood for visuo-motor paradigms^8, 11, 12^. Building on evidence that training can amplify the heritability of behavior, an intriguing possibility is that the brainprint similarity of MZ twins may be strengthened by training, whereas it may be attenuated following training in DZ twins^26, 27, 48^.

## Results

We analyzed data from 71 MZ and 34 DZ twin pairs (age 16–78 years, mean age 37 years). For each twin, we created a cortical brainprint from 272 anatomical features from T1w and T2w MRIs (Fig 1). Using this holistic metric, we quantified within-person similarity across repeated scans and between-person similarity within twin pairs, and tested associations with zygosity and within-pair differences in environmental variables.

### Brainprint similarity varies as a function of genetics and prenatal environment

Using the holistic brainprints, we confirmed that cortical anatomy acts as a robust individual signature, with perfect cross-session identification of individuals across 10 weeks (Fig. 1A). Brainprint similarity also enabled identification of MZ co-twins, who showed substantially higher similarity than DZ twin pairs (Fig. 1A).

Beyond zygosity, to identify which within-pair differences in environmental contributions were most relevant for brainprint similarity, we included for analyses a series of lifestyle-related variables. In addition to birth weight, for which we observed within-pair variance also in MZ (Table 1), the variables included current weight, BMI, blood pressure, alcohol consumption, smoking history, sleep quality^43^, physical activity^44^ and education level. Ten within-pair difference scores, along with zygosity, were entered into feature selection for brainprint similarity analyses using the random forest algorithm *Boruta*^49^. Feature selection identified zygosity and birth weight discordance as the only potentially important predictors in the full sample, whereas within the MZ sample birth weight discordance was the only retained predictor (Fig. 2A; Fig. S3). We therefore focused subsequent analyses on zygosity and birth weight discordance and tested how they associated and interacted with brainprint similarity. Main model specifications are listed in Table 3, with additional models specified in Table S1. Key results and coefficient estimates are presented in Table S2.

**Figure 2.**
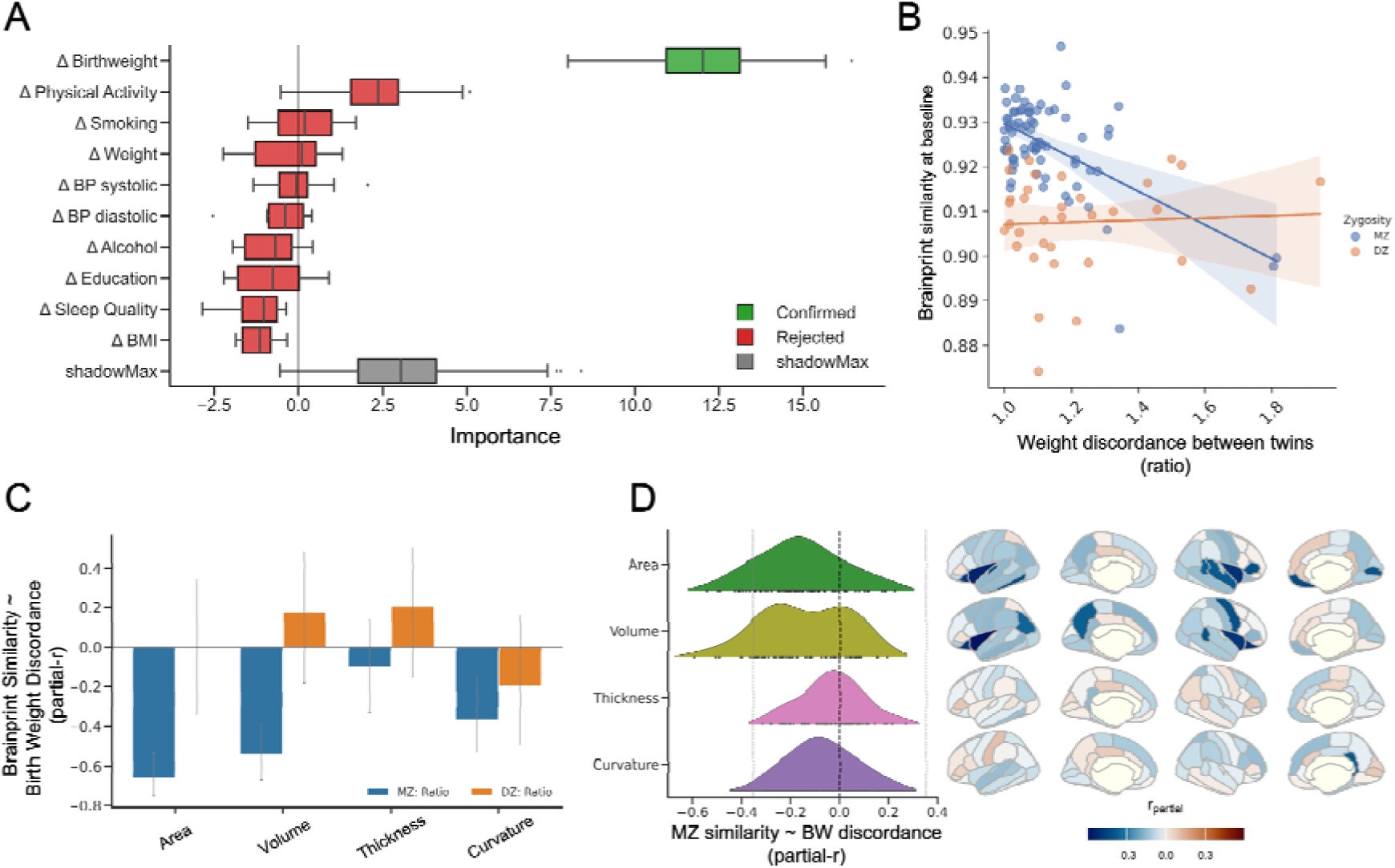
Similarity of anatomical brainprints is shaped by birthweight discordance in MZ twins. A) Feature selection using Random Forest (Boruta^49^) indicated that, among the tested within-pair environmental difference measures (Table 1), only birth weight (BW) discordance was potentially informative for brainprint similarity in monozygotic (MZ) twin pairs. In the full sample (MZ+DZ), zygosity and BW discordance were retained, whereas within the MZ sample BW discordance was the only retained predictor (Fig. S3). Boxplots show the distribution of importance scores across Boruta iterations; shadowMax (grey) is the maximum importance among permuted “shadow” features and serves as the reference threshold. Predictors exceeding shadowMax are classified as Confirmed, while those below are Rejected. B) Brainprint similarity between twins at baseline decreased with increasing BW discordance in 71 MZ pairs, but not in 34 DZ pairs (Fig. 2B). The primary test was a generalized additive model including BW discordance, zygosity, and their interaction (adjusting for age (smooth), sex, and mean BW; Table S1: Model 1), which showed a significant BW discordance × zygosity interaction (t = 4.01, p = .0001; Table S2: Model 1). For effect-size visualization, Pearson correlations are shown for MZ (r = −0.57, p < 0.0001) and DZ (r = 0.05). See Fig. S4 for alternative operationalizations of BW discordance. C) Decomposing the brainprint into feature categories showed that the BW discordance x zygosity interaction was driven primarily by cortical area and corresponding volume (t = - 7.17 and t = -5.32), with a smaller contribution from curvature (t = -3.2), while associations with thickness were not significant (Table S2: Model 1). In MZ twins, greater BW discordance was associated with lower similarity for these feature categories, whereas no corresponding associations were observed in DZ twins. Error bars reflect 95% CI. D) Feature-wise associations with BW discordance in MZ twins. Nineteen cortical features showed significant relationships with BW discordance after FDR correction (pFDR < 0.05), including ten area features and eight volume features, with one curvature feature and none for thickness. Black dots represent cortical regions; the light grey dotted line indicates the lowest partial correlation surviving FDR correction. Distributions are also plotted on brain surfaces. The scale represents the partial correlation coefficient (r-partial) from the GAM (Table S1: Model 1). Opaque regions are significant after FDR correction. Rows correspond to Area, Volume, Thickness, and Curvature. Brain views show left and right hemispheres, lateral and medial surfaces.

**Table 1.** Pair means and differences for potential predictor variables.

| Measure | MZ pairs |  |  | DZ pairs |  |  |
| --- | --- | --- | --- | --- | --- | --- |
|  | Mean | Mean | Difference | Mean | Mean | Difference |
|  |  | difference | range |  | difference | range |
| Birth weight | 2534 | 260 | 0-1490 | 2669 | 449 | 0-1750 |
| Current weight | 71.4 | 5.2 | 0-25.9 | 70.2 | 11.5 | 0.1-52.6 |
| BMI | 24.5 | 1.8 | 0-10.2 | 23.9 | 3.9 | 0.1-17.8 |
| BP systolic | 123 | 8.82 | 0.2-53.7 | 117 | 9.0 | 0.3-33.5 |
| BP diastolic | 78.2 | 7.2 | 0.5-36.3 | 75.0 | 7.0 | 0.3-21.3 |
| Alcohol Units/week | 4.3 | 3.29 | 0-32 | 2.9 | 4.82 | 0-16 |
| Smoking history | 0.2 | 0.2 | 0-2 | 0.3 | 0.4 | 0-1 |
| Education | 15.7 | 0.9 | 0-6 | 15.9 | 1.7 | 0-5 |
| Sleep quality | 6.0 | 2.1 | 0-7 | 5.9 | 2.8 | 0-8 |
| Physical activity | 1829 | 1494 | 0-6792 | 2192 | 1857 | 15-7816 |

*Age and education are given in years, birth weight in grams, current weight in kilograms, blood pressure (BP) is the average of two measurements, smoking history was indexed as 0 = never smoked, 1 = previously smoked, 2 = currently smoking. The Sleep quality scale maximum = 21, with higher values indexing worse quality<sup>43</sup>. Physical activity is given in minutes reported per week<sup>44</sup>.*

We first evaluated how baseline brainprint similarity was associated with prenatal environmental variance, indexed by birth weight (BW) discordance across all 105 twin pairs. Brainprint similarity decreased with increasing BW discordance, but this effect depended on zygosity: greater BW discordance was associated with lower brainprint similarity in MZ twins, whereas no corresponding association was observed in DZ twins (Fig. 2B; Table S2: Model 1).

Decomposing the brainprint into specific metrics indicated that the BW discordance effect in MZ twins was driven primarily by cortical area, with additional contributions from volume and curvature, whereas associations with thickness were not significant (Fig. 2C). In DZ twins, BW discordance was not significantly associated with any cortical feature category (Fig. 2C).

To identify the anatomical loci underlying the global brainprint difference effect, we mapped feature-wise associations with BW discordance in MZ twins (Fig. 2D). Nineteen cortical features showed significant relationships with BW discordance after FDR correction, with effects concentrated in area and volume measures. Across significant features, greater BW discordance was consistently associated with lower brainprint similarity, with associations distributed across frontal, parietal and temporal regions as well as the insula and cingulate cortex (Fig. 2D).

### Nature of birth weight associations

Given that BW discordance effects on brainprint similarity were driven primarily by cortical area, we next examined whether lower BW is associated with smaller cortical surface area^21–23, 35, 36^. Across the full sample, BW was positively associated with cortical area after adjusting for age and sex (Table S2: Model 2), and this association remained significant when additionally controlling for intracranial volume (ICV), suggesting an effect on cortical surface expansion beyond global head size scaling (Table S2: Model 3). It has previously been questioned whether BW correlates within MZ twins differ from those observed within DZ twins^36^. When restricting the analysis to the MZ sample, the BW-area association was stronger, consistent with the current zygosity-dependent BW effects observed for brainprint similarity (Table S2: Model 2). Importantly, the association remained robust after accounting for current body mass index (Table S2: Model 4).

Because cortical area is traditionally measured at the white matter surface, we also tested whether the BW-area association was evident across the cortical mantle by repeating the analysis using cortical area measured at the pial surface. This confirmed an equivalent positive association between birth weight and gray matter surface area, and this association was also apparent within the MZ subsample (Table S2: Model 2).

### Cortical changes in response to navigation training in virtual reality

Specific, intensive environmental demands may induce cortical changes in adulthood, with potentially different effects on brainprint similarity in MZ and DZ twins. To test this possibility, we administered a targeted 10-week spatial navigation training program in the virtual town Plasti City using immersive virtual reality (Fig 3A; see Methods). Longitudinal changes in brainprint similarity were modeled across three time-points: baseline, immediately post-training, and at follow-up (∼10 weeks post-training; Table S1: Model 5).

**Figure 3.**
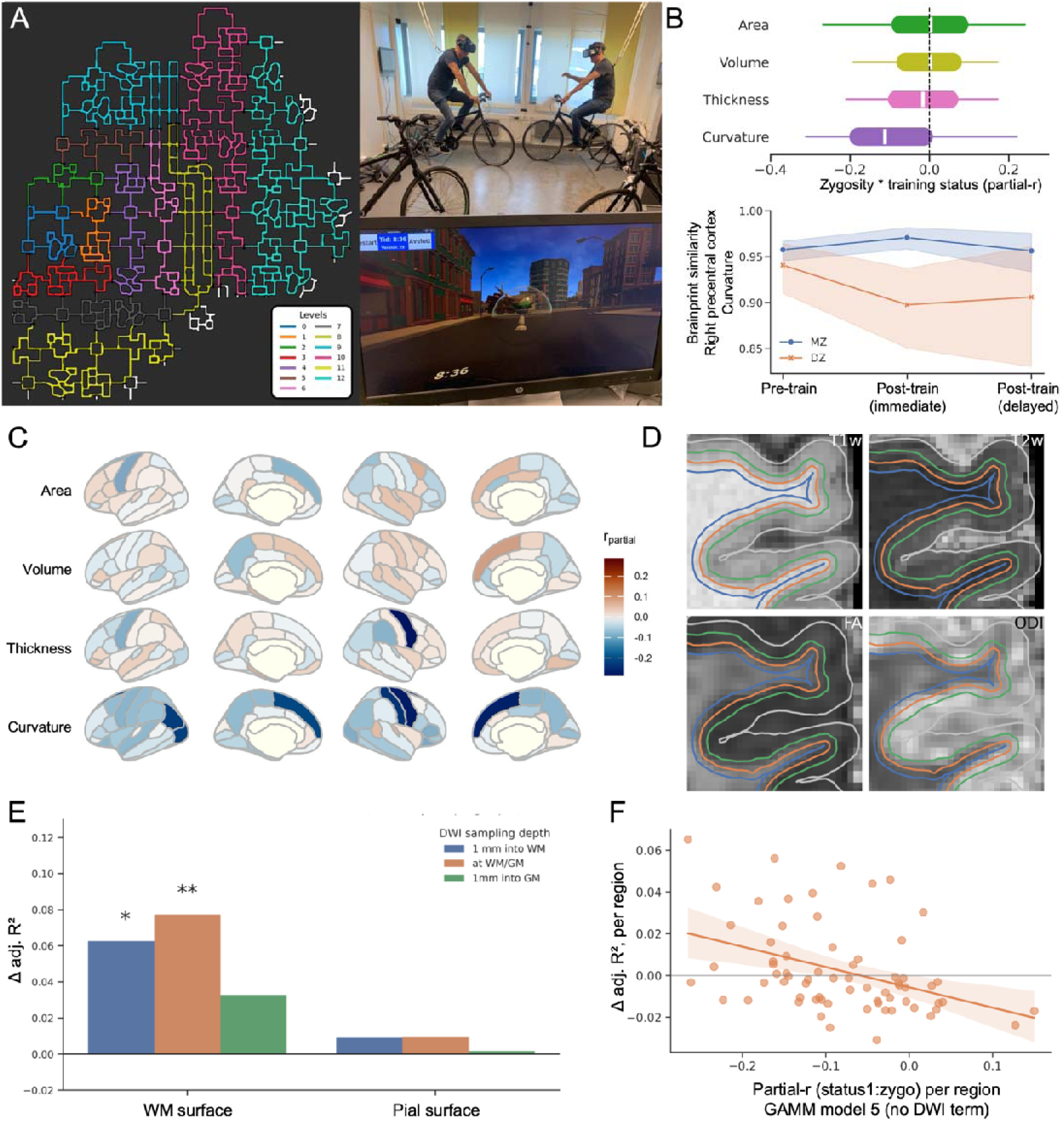
Immersive training influences anatomical and microstructural brainprint similarity. A) The virtual environment and experimental setup. Left panel: Bird’s-eye view of “Plasti City”, roads networks are colored by difficulty levels, which were opened successively according to participants’ performance. Right panel: the cybercycle set-up. The assistant screen shows what a participant sees in VR. A target is shown above the bicycle handlebars, with elapsed time in the session. In the homework version, the same is shown on the laptop screen. B–C) The effect of training status and zygosity on brainprint similarity. Restricting the analyses to valid trainers with complete pre- and post-training MRI (30 MZ and 14 DZ pairs), global brainprint similarity showed a significant training x zygosity interaction immediately post-training (t = -2.324, p = .022), which was attenuated at follow-up (t = - 1.287, p = .20) (Table S2: Model 5). B) Top: distributions of feature-wise interaction effects (r-partial) by feature dimension. The interaction was stronger for curvature than for area, volume, or thickness (Kruskal–Wallis H = 37.72, p < .0001; Dunn’s post-hoc, Bonferroni-corrected p < .0001 for curvature versus each other category). Boxplots show the median (white line), interquartile range (colored box), and whiskers extending to 1.5 x IQR. Bottom: example interaction from the region with the strongest curvature effect (right precentral cortex), illustrating increased similarity in MZ twins relative to DZ twins following training. C) Feature-wise interaction effects across regions in the full longitudinal sample including non-trainers. Surface maps show r-partial for the training status (immediate post-training) x zygosity interaction for each feature dimension (Area, Volume, Thickness, Curvature). Negative values indicate that MZ twins became more similar with training relative to DZ twins. Opaque regions indicate pFDR < .05; effects were primarily observed for curvature features, with one additional effect in right precentral thickness. D-F) Improvement in model fit when including “diffusion brainprints” in predicting cortical curvature similarity. For each session and twin pair, global microstructural twin similarity was computed from individual diffusion brainprints (6 metrics x 68 parcels), sampled at three depths relative to the WM/GM interface. D) Sampling surfaces visualized as colored outlines 1 mm into WM (blue), at the WM/GM interface (orange), and 1 mm into GM (green), together with the pial surface (gray) shown for reference. The WM/GM interface is identified in the T1w image and then expanded or contracted by 1 mm. The surfaces were projected onto the co-registered T2w image, and DTI and NODDI maps, where FA and ODI are shown as examples. E) Adding diffusion similarity and its interaction with training to the curvature-similarity model (Table S1; GAMM model 8) increased adjusted R2 relative to the baseline model that did not include diffusion terms (GAMM model 5) when curvature was measured at the WM surface (left) but not at the pial surface (right). The gain in explanatory power was largest at the WM/GM interface (adjusted ΔR² = 7.7%, pFDR < 0.004) and 1 mm into white matter (adjusted ΔR² = 6.3%, pFDR < 0.027), whereas sampling 1 mm into gray matter yielded only a weak, non-significant trend (adjusted ΔR² = 3.2%, pFDR = 0.15). Model comparison via likelihood ratio tests, FDR across depths within surface, alpha = .025 per surface; * p < .05, ** p < .01 (see Table S2: model 8) F) Regional analysis: observed improvement in model fit (y-axis) versus the strength of the training status x zygosity interaction documented above (see plot B). Parcels with stronger immediate training-related effects on curvature similarity also showed larger improvements in model fit when adding local diffusion similarity (Spearman’s rho = -0.46, p < 0.0001), whereas no such spatial correlation was observed with pial curvature (Spearman’s rho = - 0.02). Negative x-axis values indicate MZ > DZ similarity change following training. In the current plot, diffusion was sampled at the WM/GM interface and curvature estimated at the WM surface.

To provide a strict test of training-related change, we first restricted analyses to twin pairs with valid training engagement and complete pre- and post-training MRI (30 MZ and 14 DZ pairs). Using the full brainprint, we observed a significant training x zygosity interaction immediately post-training (t = -2.324, p = .022), indicating that MZ twins became more similar relative to DZ twins after training (Table S2: Model 5). This effect was attenuated at follow-up (t = -1.287, p = 0.20). Comparing brainprint feature categories, the interaction was significantly stronger along the mean curvature dimension compared to area, volume, or thickness (Fig. 3B, top). To illustrate the effect profile, we show the interaction for the region with the strongest curvature effect (right precentral cortex; Fig. 3B, bottom).

We investigated the generalizability of training-related effects detected in the curvature brainprints across the cortical mantle (30 MZ and 14 DZ pairs; no non-training). The training status x zygosity interaction in curvature similarity was present for measures defined at the white matter surface (t = -3.312, p = .0012), whereas no corresponding effect was observed for pial-surface curvature similarity (t = 1.313, p = .1915). We next tested whether training-related curvature similarity effects at the white matter surface were accompanied by directional changes in the underlying features. Specifically, we ran GAMMs predicting global mean curvature (and, for comparison, total cortical area) as a function of training status (for full model specifications see Table S1: Model 6-7). We observed no significant main effects of training status on global mean curvature at the white matter surface and no corresponding effects on pial-surface curvature, nor on global area at either surface (Table S2: Model 6-7).

### Behavioral relations of brainprint changes

Given the zygosity-dependent training effects on brainprint similarity, it is important to ascertain that these were not driven by systematic differences in training engagement between MZ and DZ pairs. Within-pair discordance in total training time did not differ significantly between MZ and DZ groups (Mann-Whitney U = 160, p = .212). However, within-pair differences in training performance (level reached in the navigation task) were smaller for MZ than DZ twins, consistent with genetic constraints on performance (Mann-Whitney U = 57, p < .0001, mean difference: MZ = 1.10, DZ = 3.71). Moreover, we observed evidence consistent with developmental contributions to training performance: in 41 MZ twin pairs with valid training data (independent of post-training MRI status), birthweight ratio (heaviest/lightest co-twin) correlated positively with difference in level reached (Pearson r = .347, p = .028), and there was a tendency for difference in training time with the heavier twin training more, albeit non-significant (r= .286, p=.073). We therefore tested training engagement and performance more formally by including within-pair differences in training time and performance as covariates and interaction terms in longitudinal models of brainprint similarity (Table S1: Model 9-10).

First, we modeled brainprint similarity as a function of zygosity and training while accounting for within-pair differences in total training time. Training time discordance interacted with training status such that greater within-pair differences in training time were associated with lower brainprint similarity immediately post-training. Put simply, brainprints turned less similar if one twin trained more than the other. Importantly, the main zygosity-dependent training effect (as visualized in Figure 3B) was preserved when controlling for training time discordance, also when restricting similarity estimation to cortical curvature features only, and when focusing on the subset of features showing FDR-corrected significant training × zygosity effects in the main analysis (Fig. 3C; Table S2: Model 9).

Second, we examined brainprint similarity in relation to within-pair differences in navigation performance (level reached after training). Here, performance discordance interacted with training status immediately post-training, indicating that twins who achieved more similar performance levels also showed higher brainprint similarity. This performance metric, which is itself genetically and developmentally constrained (as shown above), absorbed much variance, rendering the main zygosity-dependent training effect not significant in this model (Table S2: Model 10). Together, these analyses link post-training brainprint similarity to training engagement and behavioral performance, while indicating that zygosity-dependent effects are not explained by training-time discordance between MZ and DZ pairs.

### Distinct cortical feature dimensions reflect distinct environmental influences

A direct comparison of the anatomical targets revealed an apparent dissociation between the identified early-life and adult environmental influences when evaluated within the same sample and analytical framework. Using region-wise brainprint similarity models with a zygosity x environment interaction term, prenatal variance (indexed by birthweight discordance) mapped predominantly to cortical surface area (and consequently volume), whereas adult navigation training effects were manifested chiefly in cortical curvature (with more limited effects on thickness). Figure 4 visualizes this contrast directly by showing the distribution of region-wise zygosity x environment interaction effects (partial-r) across the same cortical feature dimensions and the same 105 twin-pair sample for BW discordance (panel A) and training status (panel B; corresponding to the regional effects underlying Fig. 3C). This comparison demonstrates a degree of specificity to human cortical plasticity: while cortical surface area is associated with birthweight as a proxy of intrauterine environment, adult training effects are here most evident at the gray-white matter boundary, reflected in curvature-based similarity and its coupling to microstructural brainprints.

**Figure 4.**
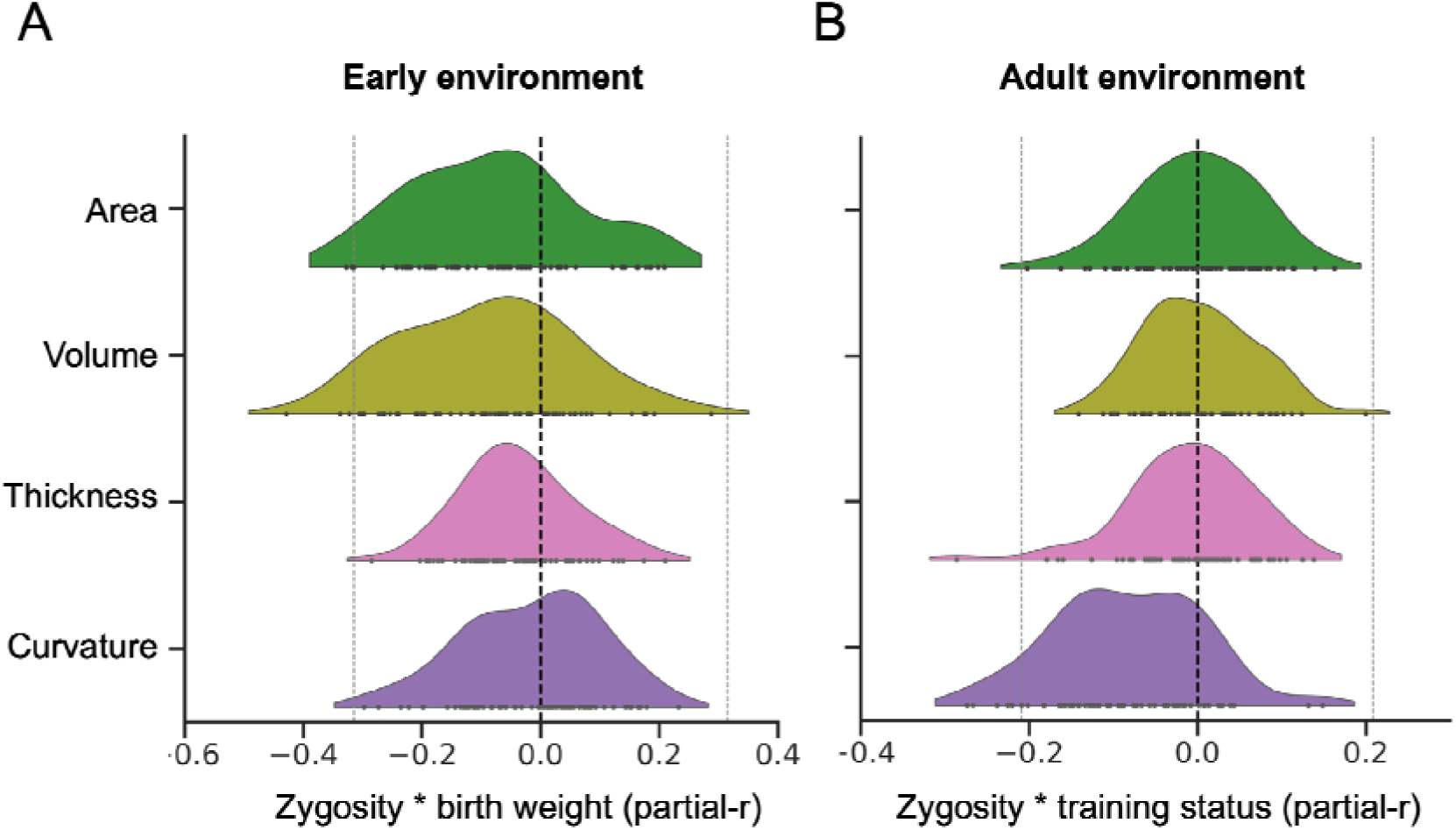
Differing distributions of gene-environment associations across cortical feature dimensions for early versus adult environmental influences. A) Early environment, prenatal variation: region-wise zygosity x birth weight discordance interaction effects on brainprint similarity (partial-r) estimated from the full twin sample (105 pairs), evaluated feature-by-feature across regions. Differences across feature categories were assessed by comparing the distributions of region-wise partial-r values using a Kruskal–Wallis test (H = 13.56, p = .0036). Dunn’s post-hoc tests (Bonferroni-corrected, only showing significant associations) indicated differences between Area and Curvature (p = .0262) and Volume and Curvature (p = .0078). B) Adult environment, training-related variation: region-wise zygosity x training status interaction effects on brainprint similarity (partial-r) estimated from the same full sample; values correspond to the regional effects underlying Fig. 3C and are expressed as partial-r to enable comparison across models. Kruskal-Wallis H = 41.32, p < .0001; Dunn’s post-hoc indicated differences between Area and Curvature (p = < .0001), Volume and Curvature (p < .0001), and Thickness and Curvature (p = <.0001). Individual datapoints represent partial correlation coefficients for individual cortical regions; the light grey dotted line indicates the lowest partial-r surviving FDR correction across the 272 region x feature tests within each panel.

## Discussion

Utilizing a combined observational and experimental twin design, the results demonstrate that genetic and environmental influences on human cortical anatomy may be teased apart. While differences in early prenatal growth were related to lasting deviations in cortical similarity in genetically identical twins, their cortical anatomy stayed similar, or even converged, with later training, relative to that of fraternal twins, which diverged. We also observe a dissociation in how these environmental influences interact with genetic factors to affect specific cortical characteristics, suggesting that different cortical features vary in their susceptibility to change depending on the type of environmental influences.

The findings show a persistent effect of prenatal environmental influences on cortical area, as seen both at the white-gray matter interface and at the pial surface. Our results also indicate that while prenatal environmental differences had strong effects on adult cortical anatomy, present variation in adult life-style-related factors among genetically identical twins was not of importance for current brainprint similarity. For instance, while MZ twins on average differed by ∼10% in birth weight, they also showed current adult weight and BMI discordances of ∼7%, but the latter differences were rejected as features of potential importance, and the association of birth weight and current cortical area was upheld when controlling for current BMI.

Cortical area and curvature, as measured at the white matter surface, still appear to be adaptable to some extent later in life, whereas such plasticity was not traceable in global measures at the pial surface. Hence, the present findings suggest that early-life influences indexed by birth weight persistently shape cortical area and the gray matter surface through life, both as main effects and in interaction with genetic factors. Still, plastic changes at the cortical white-gray interface may take place with immersive cognitive training in adulthood, here also interacting with genetic factors.

The persistent association between early environment and brainprint similarity in genetically identical twins aligns with studies linking prenatal growth to select cortical characteristics across the lifespan. We have previously shown that birth weight relates positively to adult cortical volume and area in a topographically stable way^23^. Other studies have also shown enduring effects of birth weight discordance on the brains of MZ twins^35, 36, 50^, though not invariably ^51^. For cortical morphology, two studies reported effects of birth weight discordance on cortical area and volume, though one study was restricted to MZ twins^35^, whereas the other found no interaction with zygosity^36^, thus concluding that birth weight correlates in DZ twins were not significantly different from those observed in MZ twins. In contrast, we find that associations with cortical characteristics are substantially stronger for birth weight variations in MZ twins. This can, but need not, signify other, or non-dimensional mechanisms by which environment works on birth weight and cortical features in MZ. As the genetic variance in samples of MZ twins will only be across-pair, but within and across-pair in samples also including DZ twins, the relative effect of environmental versus genetic differences affecting birth weight variation will be greater in MZ samples. Methodological differences then may contribute to the observed discrepancies. For instance, previous studies utilized CIVET ^35, 36^ (http://www.bic.mni.mcgill.ca/ServicesSoftware/CIVET) whereas the current study employs FreeSurfer (https://surfer.nmr.mgh.harvard.edu/) and brainprints^39, 52^. Using FreeSurfer, we previously showed an age-invariant effect of birth weight discordance on cortical area in genetically identical twins^23^. The present findings demonstrate the robustness of this environmental effect, showing that the multi-feature cortical brainprint^53^ diverges in genetically identical relative to fraternal twins. While birth weight variation across DZ twins and the general population reflects a combination of genetic and environmental factors, the variance observed within MZ twin pairs is necessarily environmental.

The enduring association of birth weight and cortical brainprint differences in MZ twins sampled across the adult lifespan corresponds with evidence that select processes in cortical development exclusively happen before birth. For instance, neurogenesis takes place almost only in fetal development, even though synaptogenesis, synaptic remodeling and myelination are protracted processes after infancy^2, 31, 54–58^. Humans are born with almost all cortical neurons they will have through life, and neuronal migration and differentiation are also defined early^2^. Factors that affect placental function and uterine or umbilical blood flow on a chronic basis may lead to restricted fetal growth, including brain growth, with stable^23, 59^ and especially strong effects on cortical area at white and pial surfaces, in accordance with the radial unit hypothesis^2^. These effects are isolated, and may be amplified in MZ twin pregnancies, but may also work dimensionally to some extent across DZ and singleton pregnancies too.

While behavioral heritability can increase with training^27, 48^, the present findings indicate that similar environments in adulthood can foster brain resemblance in genetically identical individuals, as training either preserve similarity or causes cortical changes to occur in parallel. The interaction between zygosity and training is driven not only by this, but largely by fraternal twins becoming less similar with training. Navigation training is a suitable paradigm for inducing such changes, as the ability to navigate shows wide variability relating to both genetic and environmental differences^45, 46^, with demonstrated effects on brain structure^11, 47^. The observed gene-environment interaction on cortical brainprints thus aligns with evidence from behavioral studies: Since genes work through environments, genetic effects will become more evident given similar environmental opportunity^27, 48, 60, 61^, causing genetically different individuals to become more different and genetically identical individuals to stay the same or even become more similar. The zygosity - training interaction on cortical brainprints offers a possible neural substrate for this phenomenon, and is in line with the observation that behavioral training causes a wider distribution of individual differences, even if the relative proportion explained by genetic factors increases^48^.

While both birth weight differences and training showed positive effects on cortical area and curvature, the curvature changes following training in adulthood were smaller and may partially reflect subtle reconfigurations of the underlying white matter surface. This is indicated by increase in total white matter area, but not in pial cortical area, with training, as shown in the full sample longitudinally. Curvature is associated with gyrification, and rapid growth of intracortical neuropil, addition of glial cells, and enlargement of subcortical white matter in primates, are the primary forces responsible for the post-neurogenic expansion of the cortical surface and formation of gyri during fetal development^62^. The same mechanisms cannot be assumed to underlie changes in curvature in adulthood, especially as these changes appear to occur in the absence of changes in pial curvature and area. Supplementary diffusion analyses indicated microstructural contributions to the training-related effects at the gray-white cortical interface, with patterns compatible with fiber organization changes on the white matter side and neurite-density on the gray matter side (Figure S6). Changes in curvature at the white matter surface, white matter area, and cortical thickness, including in deeper gray matter layers, could reflect shifts in intracortical myelination and synaptic remodeling during adulthood^37, 63^. Additional studies and measures are however required to resolve underlying mechanisms.

While the influences observed at prenatal stages and in adulthood are not directly comparable, the results underscore the substantial impact of fetal and early-life influences relative to variables typically measured in adulthood^25^. This is not only due to the timing of neural development, e.g. corticogenesis^2^, but also that effects of early-life environment are carried out over years and in interactions with the later environment^64^, whereas adult influences work over shorter time. This principle is demonstrated in our study: while MZ twins reached more similar levels of performance, the heavier twin tended to train more and attain a higher final level. While observed in a restricted 10-weeks experimental setting, these relations suggest how subtle early-life differences may scale into more substantial real-world effects taking place across decades.

Cortical representations, as here captured with brainprints show the extent of individuality of human brains^39, 40^, and can be used to decompose how this individuality may unfold through the life course, based on genetic and environmental variance. As pointed out by Sidman and Rakic, ontogenetic mechanisms must be comprehended in terms of the prolonged time of maturation, large size and other properties peculiar to the human brain ^65^. The long human lifespan allows a temporal resolution whereby we can truly observe the persistence of early-life environmental associations with specific cortical features, coupled with preserved plasticity of others. Further understanding of such fundamental concepts should thus come also from human studies^33, 65^.

Longitudinal combined observational and experimental designs including neuroimaging of MZ and DZ twins may be used to advance our understanding of what works when, and for how long, on the human brain. The current paradigm and results suggest a possible model for how associations between different types of environmental variation and human brain characteristics can be studied across the human lifespan. The findings indicate that, in the present twin sample, adult brain anatomy was to a major degree shaped by genetic and early environmental factors, while additional and more restricted influence by adult environmental events was also detectable. Variation in adult cortical anatomy may thus be traced to genetic, prenatal, and contemporary environmental factors, each with partly distinct brainprint signatures.

## Materials and methods

### Sample

The project was approved by the Regional Committee for Medical Research Ethics, South Norway, and all participants gave written informed consent. The full co-twin MRI sample reported on here, constitutes a subsample of the full Set-to-change study sample, where full twin pairs with MRI data for both twins, comprised 210 individuals (age range 16-78 years at baseline), of whom 142 MZ and 68 DZ twins. For this sample, 4 sets of MRI data for participants scanned at wave 1 were excluded due to scan artefacts (2) or lost due to human error (2), but these participants had MRI data at waves 2 and 3, and were included in analyses where appropriate. The majority of the present sample (n = 190; 90.5%) was recruited through the Norwegian Twin Registry (NTR) ^66^, based on availability of registry data for birth weight of both twins in pairs. Because the Norwegian Institute of Public Health, which the NTR is part of, could not resume recruiting for a prolonged period following additional task burden related to the COVID-19 pandemic, a minority of the sample (n = 20) was recruited through social media. Only same-sex pairs were recruited. For the majority of the sample (n = 190), zygosity information was retrieved from the National Twin Registry, while for ten pairs, this information was retrieved by current questionnaire assessment, and for the majority of the sample, DNA was also retrieved from either saliva or buccal swabs (n = 158), in all cases confirming the NTR zygosity information. (For details on determination of zygosity based on DNA, see SI, Study design and sample, Genotyping^67, 68^.) See Table 1 and details below for a description of the sample characteristics.

Participants were recruited to be healthy, community-dwelling volunteers, without worries about their cognitive function or diseases known to affect central nervous system functioning. Twins above 40 years of age were required to score a minimum of 25 on the Mini Mental Status Examination (MMSE). In the broader Set-to-change study, 231 individuals were included, and 223 underwent MRI. All research MRI data for one participant was unfortunately lost due to human error. A radiologist evaluated scans included for clinical purposes (see below), and data for 1 participant was excluded due to a clinical finding. For the remaining MRI sample (n = 221) there were 11 cases where valid structural MRI data were only available for one twin, and these were excluded from the present analyses. Additionally, two twin pairs (one MZ, one DZ) missed DWI data for at least one twin and were thus excluded from the DWI analyses.

### Birth weight measures

For all participants recruited through the Norwegian Twin Registry (NTR) born 1967 or later, birth weight information was obtained from the Medical Birth Registry of Norway (MBRN), which is a national health registry containing information about all births. For participants born before 1967, NTR records of self-reported birth weight were used from 1) an earlier questionnaire survey (Q1) conducted 1979–1982^69^ (n = 32, including for one participant, co-twin-report from this survey), or, 2) self-reports collected in 2014 by NTR as part of the twin study Social Factors and Health (n =2). For those not recruited through NTR, current self-report was used (n = 20). A high reliability of self-reported birth weight over time has been found in broader NTR samples^70^. Birth weight discordance of twins was calculated as a continuous variable, where the difference within-pair in grams yielded a negative value for the lighter and a positive value for the heavier twin. A ratio score for birth weight (twin/twin) was used for discordance analyses on brainprints.

### Lifestyle-related measures

Since the studied outcomes are within-pair similarity metrics, only variables for which within-pair differences could be meaningfully calculated were potential predictors. Adult weight, BMI, and blood pressure measures were obtained by research assistants at initiation of tests or training. Systolic and diastolic blood pressure was measured twice, and the average was used for present analyses. The other measures were based on self-report: Subjective sleep quality was assessed using the Pittsburgh Sleep Quality Index (PSQI) global score, which summarizes perceived sleep quality over the past month, with higher scores indicating poorer sleep quality^43^. Physical activity was assessed using the International Physical Activity Questionnaire (IPAQ) global score, which summarizes total self-reported physical activity volume over the past week across walking, moderate, and vigorous activities, with data cleaned and truncated according to IPAQ guidelines^44^. Alcohol consumption was self-reported units per week. Smoking history was indexed by reports of having never smoked (0), having previously regularly been smoking (1) or currently regularly smoking (2) cigarettes. Education was indexed by reported number of years to highest degree held. All twin pairs had complete records for birth weight, current weight, BMI, blood pressure (BP, systolic and diastolic) and education. Missing values within pairs prevented calculation of difference scores for physical activity level and smoking history in 1 MZ pair, and for sleep quality in 6 MZ pairs – these values were replaced by sample means for the random forest feature selection analyses.

### The intervention

Participants partook in follow-up assessments, as well as a navigational training intervention, being immersed in virtual reality (VR) in our lab and on laptops at home, searching for target objects and performing retrospective and prospective memory and time tracking tasks in a virtual town called Plasti City. Plasti City covers approximately 5 km^2^, and sequential segments (levels, 0-12) were opened automatically according to participant navigation performance. Illustrations of the city with the different levels and basic navigation task are shown in Figure 3A, and the distribution of all navigation targets is shown in Figure S7. In addition to navigating to targets, participants got various retrospective, prospective and time tracking tasks as the training progressed (see SI, Cybercycle training program.). Twins were randomly assigned to start with either a training (A) or rest (B) condition. After 10 weeks, repeated scans and cognitive assessments were undertaken, and participants were to switch conditions (A-B/B-A). After another 10 weeks, scans and cognitive assessments were again repeated (see Table 2). Similarly, participants not training were invited for repeated scans and cognitive assessments at 10 week intervals. While the intention was to schedule assessments within 11-12 week intervals, due to COVID-19-related lockdown and prolonged restrictions and practical difficulties following these (see SI, Study design and sample, Data collection), the actual intervals varied more. Mean interval between wave 1 and 2 assessment in the training groups was 0.21 years (SD: 0.03, range: 0.16-0.37 years), and mean interval between wave 1 and 3 assessment was 0.44 years (SD: 0.07, range: 0.22-0.81 years). Mean interval between pre- and post-train visits was 0.21 yrs (range 0.18-0.37 years), while mean interval between post-train and post-rest-after-training visits was 0.24 yrs (range 0.18-0.58). In cases of delays in scan times, participants training were encouraged to keep training.

**Table 2.**
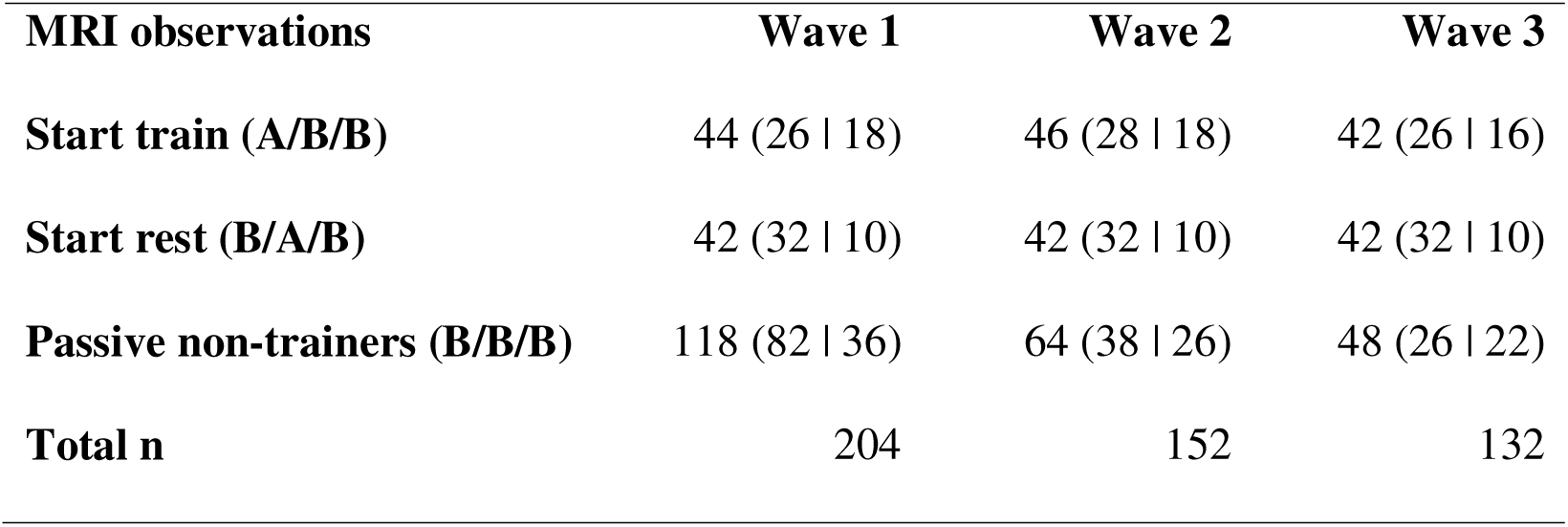
Distribution of MRI observations (n = 488) for trainers and passive non-trainers across waves. Numbers indicate observations/participants separated into (MZ | DZ) inside parentheses. MRIs immediately post-training period (n = 88) are denoted by Start train observations for wave 2 and Start rest observations for wave 3. There are 42 additional observations after ending training and resting for ∼10 weeks, denoted by Start train observations for wave 3. Note that these numbers reflect the final trial classifications, where 9 twin pairs initially enrolled in training were classified as passive non-trainers because they did not meet the minimum engagement criteria (≥2 hours training and progression beyond the baseline level). Sensitivity analyses with regard to such classification was performed, see results and SI.

### Behavioral data from the intervention

For the remaining 88 participants with more than two hours of training and MRIs post-training, mean training time was 15.18 hours, but time spent training varied much, ranging from 2.24 to 55.53 hours (SD: 11.83 hours). An average of 5.95 hours of the training time was spent in immersive VR (SD: 3.08 hours; range: 0.28 to 13.56 hours). On average, level reached was 6.64 (range: 1-12, SD: 3.40), and mean number of targets reached was 384 (range 74-932, SD: 182). Total time spent training correlated positively with number of targets reached (r = .86, p < .0001), and with level reached (r = .56, p <.0001), but not with mean deviance from shortest path (r = .12, p = .2561). The latter may be due to older participants spending more time training (correlation of visit age and training time, r = .65, p <.0001) but still having greater mean deviance from shortest path (r = .59, p <.0001). The deviance from shortest path is calculated as the difference between the participant’s path and the shortest path, divided by the shortest path length. For additional details, see SI, Study design and sample, Data collection and Cybercycle training program. Due to the variation, brainprint analyses including training times as well as level reached as dimensional variables were also conducted.

### MRI acquisition and post-processing

Imaging data was acquired using a 32-channel head coil on a 3T MRI scanner (Prisma, Siemens, Germany) at Rikshospitalet, Oslo University Hospital. Relevant for brainprint analyses were: 1) T1-weighted MPRAGE with 208 sagittal slices (TR = 2400 ms, TE = 2.22 ms, TI = 1000 ms, flip angle = 8°, matrix = 300×320×208, voxel size = 0.8×0.8×0.8 mm, FOV = 240×256 mm, iPAT = 2); 2) T2-weighted SPACE images with 208 sagittal slices (TR = 3200 ms, TE = 5.63 ms, matrix = 320×300×208, voxel size = 0.8×0.8×0.8 mm, FOV = 256×240 mm, iPAT = 2), and 3) diffusion-weighted images (DWI) with two shells (b = 1500 and 3000 s/mm², 92–93 directions and 6 b = 0 per shell) acquired using a multi-band EPI sequence (TR = 3230 ms, TE = 89.2 ms, flip angle = 78°, FOV = 210×210 mm², voxel size = 1.5 mm isotropic, 92 axial slices, multiband factor = 4, phase partial Fourier = 6/8, PE = AP, additional b = 0 PA volumes for distortion correction, total scan time ≈ 12 min). All sequences were acquired with Prescan Normalize enabled to standardize coil sensitivity profiles. These sequences are otherwise identical to those used in the HCP phase 1b Lifespan pilot (https://www.humanconnectome.org/study-hcp-lifespan-pilot/phase1b-pilot-parameters).

Structural data from each wave were first processed using FreeSurfer’s (v.7.3.2) default cross-sectional recon-all pipeline with the optional parameters --t2pial and --hires. The --t2pial parameter uses the auxiliary T2-weighted sequence to refine pial surface placement, while --hires processes the data at its native 0.8 mm isotropic resolution for enhanced surface detail. To further optimize reconstruction, a set of “expert-options” were used. This included the critical PlaceMMPialSurf fix (--mm_min_inside 50 --mm_max_inside 200 --mm_min_outside 10 --mm_max_outside 50) to address reported pial surface misplacements, and a pial surface placement enhancement in the inferior frontal area near the putamen (WhitePreAparc --rip-bg-no-annot). Additional fine-tuning options, as outlined in the FreeSurfer v7.3.2 post-release notes https://surfer.nmr.mgh.harvard.edu/fswiki/ReleaseNotes were also incorporated.

Base templates for longitudinal processing were created using FreeSurfer’s recon-all -base command. For participants undergoing training (ABB and BAB groups), templates were constructed using imaging data from sessions immediately before and after the intervention. This approach prevented systematic biases in the template towards either the pre-training or post-training phases, ensuring a balanced representation of both phases. For non-training twins, who had no intervention, templates were created using MRI data from either sessions 1 and 2 or sessions 2 and 3, with the two variants balanced by age and sex to account for potential session-specific variability. For the subset of non-training twins with only one session, the base template was constructed from the single time point, ensuring consistent resampling and interpolation across all participants.

Finally, longitudinal processing was performed using FreeSurfer’s recon-all -long command. Each cross-sectional session was aligned to the participant-specific base template to ensure consistent registration and surface placement across time points. The T2w image was consistently included with -T2pial flags to refine pial surface placement, and all processing was conducted at the native 0.8 mm isotropic resolution via the -hires option.

DWI data were preprocessed using the QuNex container (v1.0.3)^71^ to execute the HCP Diffusion Preprocessing Pipeline (v5.0.0)^72^. In brief, after gradient-nonlinearity and susceptibility distortion correction (via TOPUP), eddy-current, motion and slice-outlier correction was performed using FSL Eddy with outlier detection and replacement enabled, flagging slices exceeding 4 standard deviations. Multiband factor was specified so that outlier replacement correctly accounted for group-acquired slices. For each session, the HCP T1w image was registered to the participant’s FreeSurfer -long space using boundary-based registration; the transform was inverted and applied to the -long white and pial surfaces so they could be used to sample diffusion data in HCP T1w space, ensuring the same FreeSurfer surfaces for anatomical and diffusion brainprints. Neurite orientation dispersion and density imaging (NODDI^73^) modeling was performed with the CUDIMOT^74^ Watson model as implemented in QuNex. Diffusion tensors were fitted in FSL DTIFIT using ordinary least squares and all non-zero b-values (b=1500 and 3000 s/mm²), following Fukutomi et al.^75^, to remain consistent with the multi-shell data used for NODDI. Mixing b-values may slightly shift absolute MD/AD/RD and FA relative to single-shell fits, however our inferences were performed on similarity measures and did not rely on normative values.

When intracranial volume was used as a covariate in analyses, this was Total Intracranial Volume from Sequence Adaptive Multimodal Segmentation (SAMSEG)^76, 77^.

### Cortical morphology measures

For the FreeSurfer brainprints, cortical area, thickness, volume, and curvature were calculated for each of the 68 cortical regions defined by the Desikan-Killiany atlas^41^. The definitions of area, thickness and volume followed FreeSurfer’s standard approach: Cortical area was measured as the total surface area at the white surface (inner boundary of the cortical gray matter), cortical thickness as the mean cortical thickness between corresponding vertices on the white and pial surfaces, and cortical volume using a surface-based method where the space between the white and pial surfaces was divided into tetrahedra for precise volume estimation.

Cortical curvature was computed as the integrated rectified mean curvature over each region, capturing the total amount of cortical folding irrespective of fold direction. While area and curvature were originally defined on the white surface, control analyses also estimated mean curvature and surface area at the pial surface to investigate whether effects observed at the white surface might reflect changes in the underlying white matter rather than cortical gray matter.

While comparable approaches using the above cortical morphology measures for individual characterization have also included FreeSurfer’s estimates of sulcal depth^78^, this measure is normalized within each session, making it a relative metric specific to individual scans^79^. As a result, sulcal depth values derived from Freesurfer are not directly comparable across time points and thus not suitable to investigate changes due to interventions, so such metrics were not included here.

### Brainprint analyses

Anatomical brainprints were constructed for each participant-session by extracting the four morphological measures of interest—cortical area, thickness, volume, and curvature—for each of the 68 regions in the DK-atlas, resulting in a vector of 272 values representing cortical morphology. Note that while the current approach is inspired by Wachinger et al.’s “BrainPrint”/shapeDNA methods^40^, we here use the term “brainprint” in a generic fashion. To compare brainprints within individuals over time or between individuals, we used the inverse of the mean relative difference as a similarity metric (Formula 1). This measure calculates the relative difference between corresponding features across brainprints, normalized by their average, and expresses the result as 1 - *Mean Relative Difference*. Values closer to 1 indicate higher similarity. The mean relative difference does not require features to be on the same scale (unlike alternative similarity measures such as cosine or Euclidean similarity), making it suitable for the heterogeneous brainprint measures. It can also be applied to reduced brainprints and even single features, enabling the targeted analyses presented here.

However, one limitation of this measure is its sensitivity to differences in brain size, particularly for features such as cortical area and volume, where individuals with similar brain sizes (e.g., same sex and age) may appear more similar. To account for this, non-twin comparisons included additional analyses with sex and age regressed out using Gaussian additive models with age as a smooth term. Full brainprint analyses were also repeated using cosine similarity as an alternative metric, which is scale-invariant and unaffected by brain size differences (see Figure S8). For twin comparisons (monozygotic and same-sex dizygotic), the mean relative difference metric was used without regressing out sex or age, as these analyses inherently controlled for these variables.

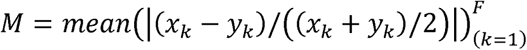

*Formula 1: Mean of the relative differences between corresponding features x_k and y_k, normalized by their average, across all features k from 1 to F. x and y can represent* brainprints *from the same individual over two different sessions or from different individuals. Brainprint similarity equals the inverse of M*.

Leveraging a set of twin pairs with two passive, no training commenced, scans (N = 30), we observed that brainprint similarity showed high precision and sensitivity to experimental manipulation (see SI and Figure S9).

### *Diffusion* brainprints

Four DTI metrics (Fractional Anisotropy, FA; Mean Diffusivity, MD; Axial Diffusivity, AD; Radial Diffusivity, RD) and two NODDI metrics (Orientation Dispersion Index, ODI; Intra-Cellular Volume Fraction – aka Neurite Density Index, ICVF), were extracted at each vertex along the Freesurfer defined WM/GM interface and averaged over DK-parcels as for the anatomical brainprints. Additional samples were taken ±1 mm along the surface normal relative to the interface (1 mm into WM and 1 mm into GM). The transition from preprocessed diffusion data in volumetric space to surface coordinates and the sampling at different depths were done using nilearn (https://github.com/nilearn/nilearn; surface.vol_to_surf)^80, 81^. Similarity of diffusion brainprints within twin pairs was estimated as for the anatomical brainprints, using the inverse of the mean relative difference of the absolute metric values.

### Statistical analyses

Statistical analyses were performed using Python 3.10.4 and R. Outside R, general statistics were computed in Python using pingouin (0.5.5). Feature selection analyses were performed using randomForest and Boruta (R 4.4. 2) and generalized additive models (GAMs) and generalized additive mixed models (GAMMs) were fitted in R 4.2.1. using mgcv and gamm4, occasionally accessed from Python via rpy2 (3.5.16).

Brainprint identification (self-identification and twin identification) was performed by ranking pairwise similarity matrices; accuracy (top-1 and top-2) and significance were evaluated against a permutation null (5000 permutations; see Fig1 caption). Feature selection over within-pair environmental difference measures was conducted using the Boruta algorithm, a random-forest-based all-relevant feature selection method algorithm in the full sample and within MZ twins. In Boruta, the importance of each predictor is assessed by comparing its random forest importance score to the distribution of importance scores obtained from permuted copies of the predictors (“shadow features”). Predictors whose importance consistently exceeds the maximum shadow importance are classified as confirmed important, while those consistently underperforming are rejected. The model was specified with baseline brainprints as the dependent variable and all remaining variables as candidate predictors. The algorithm was run until convergence, after which tentative predictors were adjudicated using the TentativeRoughFix heuristic. Only predictors classified as confirmed important after this step were selected and retained.

Three primary GA(M)M models were used for hypothesis testing (Table 3). Model 1 tested baseline brainprint similarity as a function of zygosity, birth weight discordance (and their interaction) sex, smooth age, and mean birthweight (one observation per twin pair). Model 5 tested longitudinal effects of training on brainprint similarity, including training status, zygosity (and their interaction), and smooth age, with twin pair modeled as a random effect. Model 8 extended Model 5 by additionally including diffusion-derived similarity and its interaction with training status to assess microstructural contributions to training-related brainprint changes (twin pair as random effect).

**Table 3.** Main generalized additive (mixed) models. See Table S1 for all models.

| GA(M)M | Formula | Random Effects |
| --- | --- | --- |
| 1 | Brainprint similarity ~ zygosity + sex + s(age) + mean birth weight + birth weight discordance + zygosity * birth weight discordance | NA (one observation per twin pair) |
| 5 | Brainprint similarity ~ training status + zygosity + s(age) + zygosity * training status | ~(1 \| twin pair ) |
| 8 | Brainprint similarity ~ training status + zygosity + s(age) + zygosity * training status + diffusion similarity + diffusion similarity * training status | ~(1 \| twin pair ) |

Follow-up models examining specific cortical features and robustness to alternative operationalizations are specified in Table S1 and key model coefficients are provided in Tables S2. Two-sided tests were used throughout the study, and significance was evaluated at α = 0.05.

## Acknowledgements

We sincerely thank all twins for their time and engagement in the study and for contributing to all the respective data sources. Data used were in part obtained through the Norwegian Twin Registry (NTR) and the Medical Birth Registry (MBRN) of Norway. We are grateful to NTR for recruiting the twins and providing data from their surveys as well as from MBRN. We thank the many LCBC employees who have been assisting with the training, testing, data management and processing, and Irantzu Anzar Martinez de Lagran for performing zygosity analysis. This study was funded by ERC grant 771375 and Research Council of Norway (RCN) grants to KBW. The work of JRH was partly supported by the Research Council of Norway through its Centres of Excellence funding scheme, project number 262700. The ERC and RCN had no role in the design and conduct of this specific study.

## Data Availability

The LCBC dataset has restricted access, requests can be made to the corresponding author, and some of the data can be made available given appropriate ethical and data protection approvals. However, the registry data on birth weight connected to this sample are not shareable by the authors, as these data are owned by the Medical Birth Registry of Norway https://www.fhi.no/en/hn/health-registries/medical-birth-registry-of-norway/medical-birth-registry-of-norway/, and the Norwegian Twin Registry https://www.fhi.no/en/more/health-studies/norwegian-twin-registry/ so that any access to data must be approved by them. The code used for main analyses can be accessed here https://github.com/LCBC-UiO/Set-to-change

## Author Contributions

KBW conceptualized and designed the study, did statistical analyses in R and wrote the manuscript. ACSB helped form the intervention, KEØØ did most work on designing the VR city, ACSB, KEØØ and JK helped undertake data collection, ØS and PFG conceptualized statistical models for the intervention, DVP, JK, JLAW, AMM and MS managed, post-processed and curated data, LN, NOC, JRH, YW and AMF served as consultants to study design, YW devised zygosity analysis, PDT did radiological evaluations, DVP performed reliability analysis, MHS performed brainprint imaging analyses and conducted statistical analyses, all authors commented and contributed to the manuscript draft.

## Supplementary Material

### Supporting results

**Table S1.**
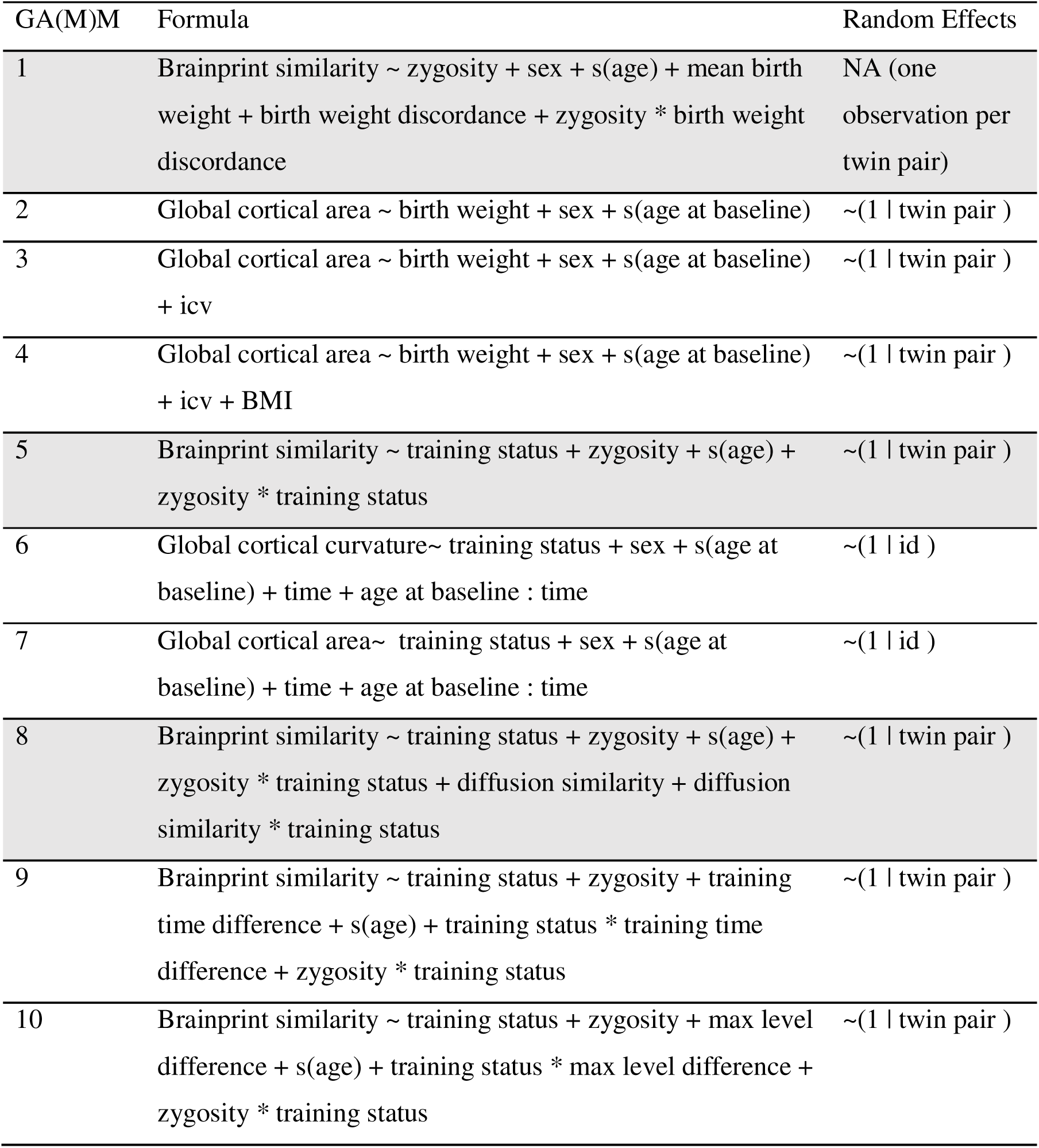
Generalized additive (mixed) models (GA(M)M) formulas run. Main analyses are shown in shaded rows, follow-up analyses in white. Explanation of terms: zygosity: ordered categorical, 1 = MZ, 2 = DZ; sex: male/female; “s(age)”: smooth-term of twin1 age in years (with 5 decimal points) at measurement; “age at baseline”: age in years (with 5 decimal points) at first measurement; “time”: time since baseline, i.e. age in years minus minimum age at measurement; “birth weight”: birth weight of single individual in grams, “mean birthweight”: (twin 1 BW + twin 2 BW)/2; “birth weight discordance”: If ratio, BW heaviest twin / BW lightest twin. If difference, BW heaviest twin – BW lightest twin; “BMI”: current body mass index; “training status”: ordered categorical, 0 = pre-training, 1 = immediately post-training, 2 = delayed post-training. “training time difference”: z-normalized absolute difference in training times between twins in a pair.

**Table S2.**
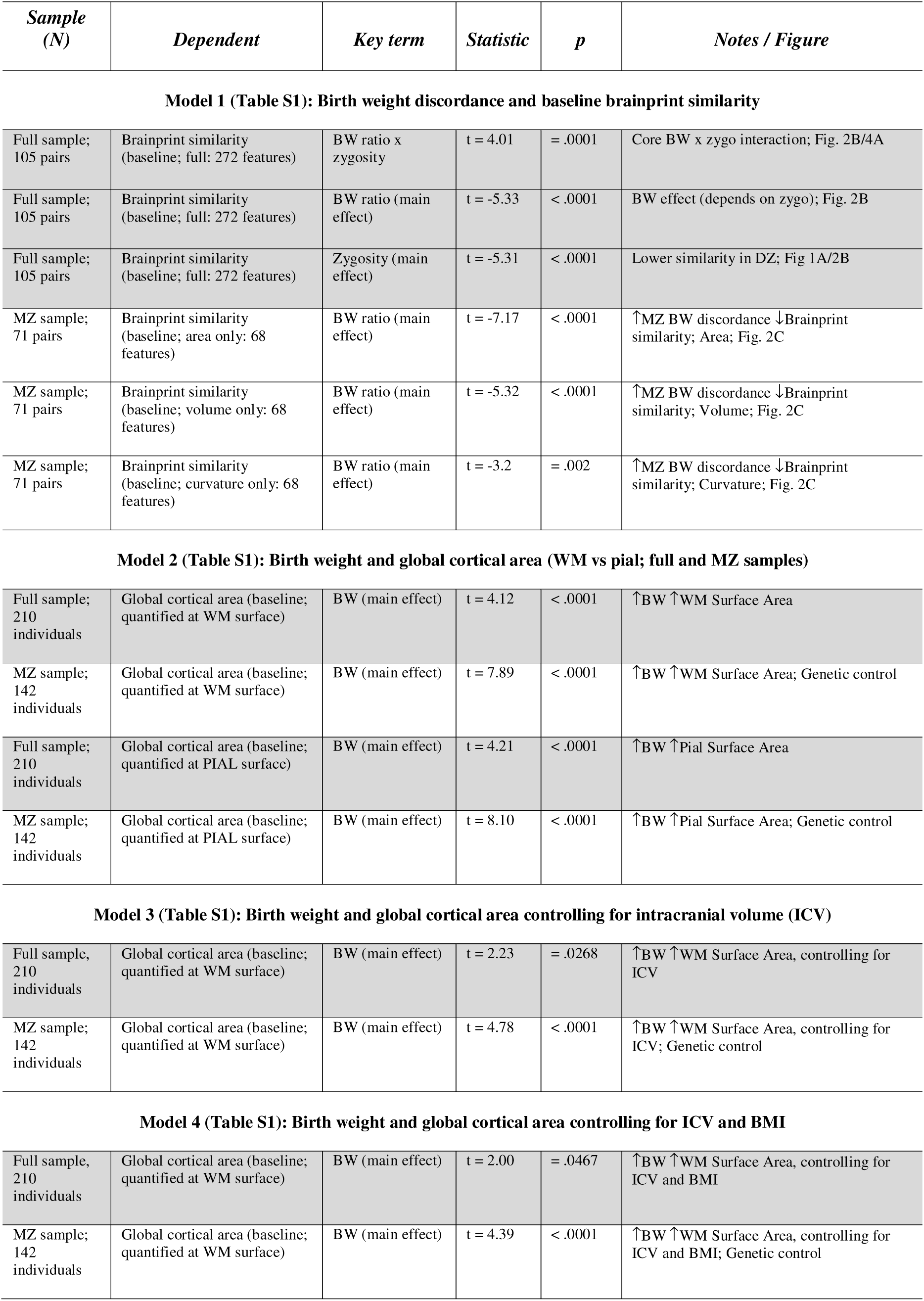
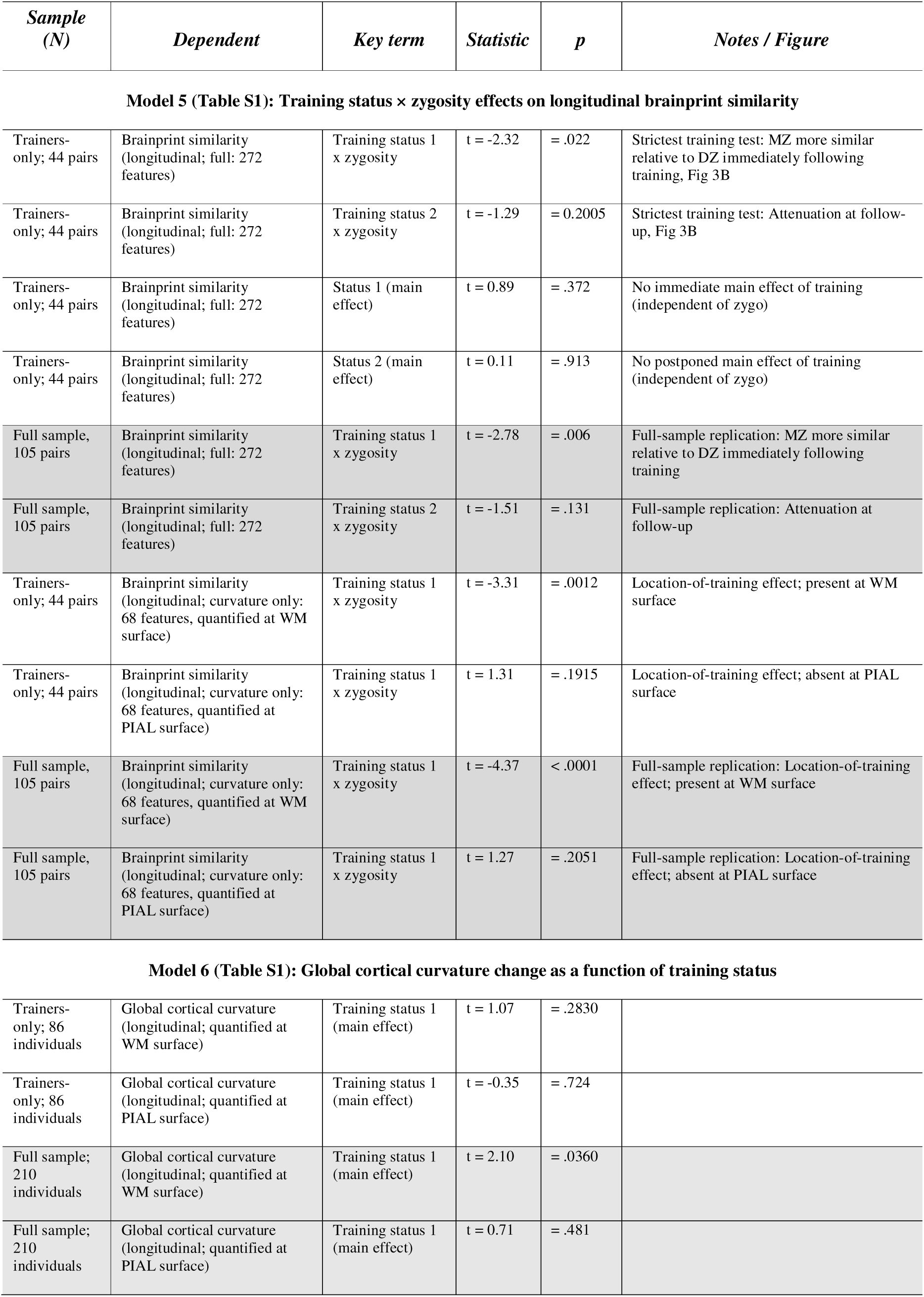
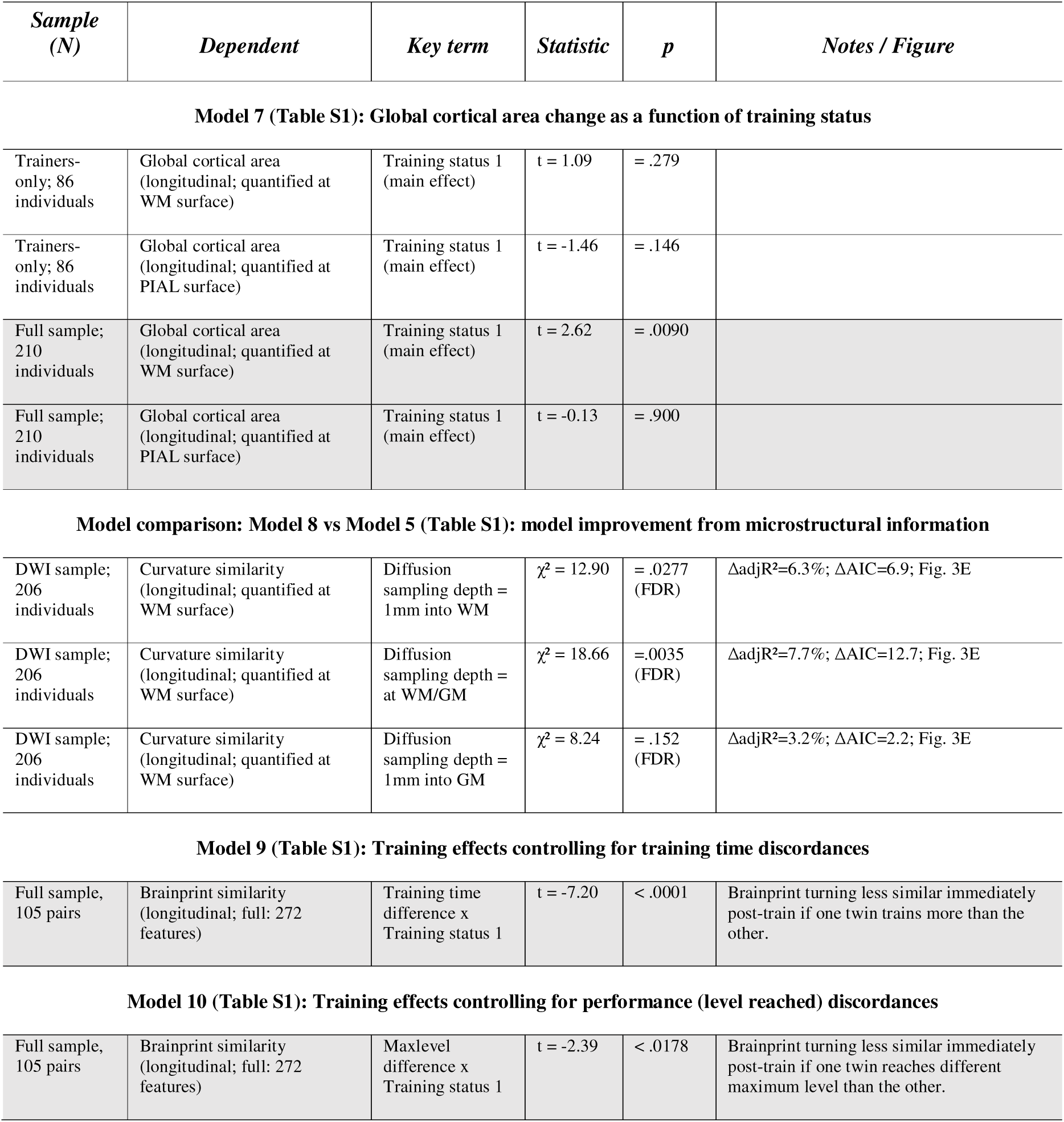
Key outputs from GA(M)M analyses. For each model specified in Table S1, the table lists the sample (pairs or individuals), dependent variable, key term(s) referenced in the main text/figures, corresponding test, and p-value. Shaded rows indicate tests involving the full twin pair sample of 105 pairs.

| *Sample (N)* | *Dependent* | *Key term* | *Statistic* | *p* | *Notes / Figure* |
| --- | --- | --- | --- | --- | --- |
| **Model 1 (Table S1): Birth weight discordance and baseline brainprint similarity** |  |  |  |  |  |
| Full sample; 105 pairs | Brainprint similarity (baseline; full: 272 features) | BW ratio x zygosity | t = 4.01 | = .0001 | Core BW x zygo interaction; Fig. 2B/4A |
| Full sample; 105 pairs | Brainprint similarity (baseline; full: 272 features) | BW ratio (main effect) | t = -5.33 | < .0001 | BW effect (depends on zygo); Fig. 2B |
| Full sample; 105 pairs | Brainprint similarity (baseline; full: 272 features) | Zygosity (main effect) | t = -5.31 | < .0001 | Lower similarity in DZ; Fig 1A/2B |
| MZ sample; 71 pairs | Brainprint similarity (baseline; area only: 68 features) | BW ratio (main effect) | t = -7.17 | < .0001 | ↑MZ BW discordance ↓Brainprint similarity; Area; Fig. 2C |
| MZ sample; 71 pairs | Brainprint similarity (baseline; volume only: 68 features) | BW ratio (main effect) | t = -5.32 | < .0001 | ↑MZ BW discordance ↓Brainprint similarity; Volume; Fig. 2C |
| MZ sample; 71 pairs | Brainprint similarity (baseline; curvature only: 68 features) | BW ratio (main effect) | t = -3.2 | = .002 | ↑MZ BW discordance ↓Brainprint similarity; Curvature; Fig. 2C |
| **Model 2 (Table S1): Birth weight and global cortical area (WM vs pial; full and MZ samples)** |  |  |  |  |  |
| Full sample; 210 individuals | Global cortical area (baseline; quantified at WM surface) | BW (main effect) | t = 4.12 | < .0001 | ↑BW ↑WM Surface Area |
| MZ sample; 142 individuals | Global cortical area (baseline; quantified at WM surface) | BW (main effect) | t = 7.89 | < .0001 | ↑BW ↑WM Surface Area; Genetic control |
| Full sample; 210 individuals | Global cortical area (baseline; quantified at PIAL surface) | BW (main effect) | t = 4.21 | < .0001 | ↑BW ↑Pial Surface Area |
| MZ sample; 142 individuals | Global cortical area (baseline; quantified at PIAL surface) | BW (main effect) | t = 8.10 | < .0001 | ↑BW ↑Pial Surface Area; Genetic control |
| **Model 3 (Table S1): Birth weight and global cortical area controlling for intracranial volume (ICV)** |  |  |  |  |  |
| Full sample; 210 individuals | Global cortical area (baseline; quantified at WM surface) | BW (main effect) | t = 2.23 | = .0268 | ↑BW ↑WM Surface Area, controlling for ICV |
| MZ sample; 142 individuals | Global cortical area (baseline; quantified at WM surface) | BW (main effect) | t = 4.78 | < .0001 | ↑BW ↑WM Surface Area, controlling for ICV; Genetic control |
| **Model 4 (Table S1): Birth weight and global cortical area controlling for ICV and BMI** |  |  |  |  |  |
| Full sample; 210 individuals | Global cortical area (baseline; quantified at WM surface) | BW (main effect) | t = 2.00 | = .0467 | ↑BW ↑WM Surface Area, controlling for ICV and BMI |
| MZ sample; 142 individuals | Global cortical area (baseline; quantified at WM surface) | BW (main effect) | t = 4.39 | < .0001 | ↑BW ↑WM Surface Area, controlling for ICV and BMI; Genetic control |

| *Sample (N)* | *Dependent* | *Key term* | *Statistic* | *p* | *Notes / Figure* |
| --- | --- | --- | --- | --- | --- |
| **Model 5 (Table S1): Training status × zygoty effects on longitudinal brainprint similarity** |  |  |  |  |  |
| Trainers-only; 44 pairs | Brainprint similarity (longitudinal; full: 272 features) | Training status 1 x zygoty | t = -2.32 | = .022 | Strictest training test: MZ more similar relative to DZ immediately following training, Fig 3B |
| Trainers-only; 44 pairs | Brainprint similarity (longitudinal; full: 272 features) | Training status 2 x zygoty | t = -1.29 | = 0.2005 | Strictest training test: Attenuation at follow-up, Fig 3B |
| Trainers-only; 44 pairs | Brainprint similarity (longitudinal; full: 272 features) | Status 1 (main effect) | t = 0.89 | = .372 | No immediate main effect of training (independent of zygo) |
| Trainers-only; 44 pairs | Brainprint similarity (longitudinal; full: 272 features) | Status 2 (main effect) | t = 0.11 | = .913 | No postponed main effect of training (independent of zygo) |
| Full sample, 105 pairs | Brainprint similarity (longitudinal; full: 272 features) | Training status 1 x zygoty | t = -2.78 | = .006 | Full-sample replication: MZ more similar relative to DZ immediately following training |
| Full sample, 105 pairs | Brainprint similarity (longitudinal; full: 272 features) | Training status 2 x zygoty | t = -1.51 | = .131 | Full-sample replication: Attenuation at follow-up |
| Trainers-only; 44 pairs | Brainprint similarity (longitudinal; curvature only: 68 features, quantified at WM surface) | Training status 1 x zygoty | t = -3.31 | = .0012 | Location-of-training effect; present at WM surface |
| Trainers-only; 44 pairs | Brainprint similarity (longitudinal; curvature only: 68 features, quantified at PIAL surface) | Training status 1 x zygoty | t = 1.31 | = .1915 | Location-of-training effect; absent at PIAL surface |
| Full sample, 105 pairs | Brainprint similarity (longitudinal; curvature only: 68 features, quantified at WM surface) | Training status 1 x zygoty | t = -4.37 | < .0001 | Full-sample replication: Location-of-training effect; present at WM surface |
| Full sample, 105 pairs | Brainprint similarity (longitudinal; curvature only: 68 features, quantified at PIAL surface) | Training status 1 x zygoty | t = 1.27 | = .2051 | Full-sample replication: Location-of-training effect; absent at PIAL surface |

**Model 6 (Table S1): Global cortical curvature change as a function of training status**
|  |  |  |  |  |
| --- | --- | --- | --- | --- |
| Trainers-only; 86 individuals | Global cortical curvature (longitudinal; quantified at WM surface) | Training status 1 (main effect) | t = 1.07 | = .2830 |
| Trainers-only; 86 individuals | Global cortical curvature (longitudinal; quantified at PIAL surface) | Training status 1 (main effect) | t = -0.35 | = .724 |
| Full sample; 210 individuals | Global cortical curvature (longitudinal; quantified at WM surface) | Training status 1 (main effect) | t = 2.10 | = .0360 |
| Full sample; 210 individuals | Global cortical curvature (longitudinal; quantified at PIAL surface) | Training status 1 (main effect) | t = 0.71 | = .481 |

| *Sample (N)* | *Dependent* | *Key term* | *Statistic* | *p* | *Notes / Figure* |
| --- | --- | --- | --- | --- | --- |

**Model 7 (Table S1): Global cortical area change as a function of training status**
|  |  |  |  |  |
| --- | --- | --- | --- | --- |
| Trainers-only; 86 individuals | Global cortical area (longitudinal; quantified at WM surface) | Training status 1 (main effect) | t = 1.09 | = .279 |
| Trainers-only; 86 individuals | Global cortical area (longitudinal; quantified at PIAL surface) | Training status 1 (main effect) | t = -1.46 | = .146 |
| Full sample; 210 individuals | Global cortical area (longitudinal; quantified at WM surface) | Training status 1 (main effect) | t = 2.62 | = .0090 |
| Full sample; 210 individuals | Global cortical area (longitudinal; quantified at PIAL surface) | Training status 1 (main effect) | t = -0.13 | = .900 |
| **Model comparison: Model 8 vs Model 5 (Table S1): model improvement from microstructural information** |  |  |  |  |  |
| DWI sample; 206 individuals | Curvature similarity (longitudinal; quantified at WM surface) | Diffusion sampling depth = 1mm into WM | $\chi^2 = 12.90$ | = .0277 (FDR) | $\Delta\text{adjR}^2=6.3\%$; $\Delta\text{AIC}=6.9$; Fig. 3E |
| DWI sample; 206 individuals | Curvature similarity (longitudinal; quantified at WM surface) | Diffusion sampling depth = at WM/GM | $\chi^2 = 18.66$ | = .0035 (FDR) | $\Delta\text{adjR}^2=7.7\%$; $\Delta\text{AIC}=12.7$; Fig. 3E |
| DWI sample; 206 individuals | Curvature similarity (longitudinal; quantified at WM surface) | Diffusion sampling depth = 1mm into GM | $\chi^2 = 8.24$ | = .152 (FDR) | $\Delta\text{adjR}^2=3.2\%$; $\Delta\text{AIC}=2.2$; Fig. 3E |
| **Model 9 (Table S1): Training effects controlling for training time discordances** |  |  |  |  |  |
| Full sample, 105 pairs | Brainprint similarity (longitudinal; full: 272 features) | Training time difference x Training status 1 | t = -7.20 | < .0001 | Brainprint turning less similar immediately post-train if one twin trains more than the other. |

**Table S2.** Key outputs from GA(M)M analyses.
|  |  |  |  |  |  |
| --- | --- | --- | --- | --- | --- |
| Full sample, 105 pairs | Brainprint similarity (longitudinal; full: 272 features) | Maxlevel difference x Training status 1 | t = -2.39 | < .0178 | Brainprint turning less similar immediately post-train if one twin reaches different maximum level than the other. |

### Supporting methods

#### Study design and sample

##### Genotyping

Briefly, the DNA samples were genotyped by the Global Screening Array (Illumina, CA). The genotypes were quality checked and imputed to the 1000 Genomes EUR population using our previous established protocol^67^. After further quality check of imputed genotypes, the PLINK2 software^68^ was used to compute the kinship coefficients based on autosomal variants which have a minor allele frequency >0.1 and are not correlated with each other (r < 0.1). Monozygotic twins were defined as those whose kinship coefficients > 0.4 and dizygotic twins are > 0.2 and < 0.3.

##### Data collection

The data collection for this study started in 2019 and ran throughout 2023, the prolonged period being due to corona restrictions during part of the project. The reasons for dropping out of training in the VR lab, MRI scanning, or project participation overall, thus may represent a mix of direct physical hindrance, and individual motivation and anxiety, the relative portions of which we unfortunately cannot meaningfully calculate, as differing restrictions and advices applied per participant timing, address, age group and health status. Hypotheses for an fMRI paradigm for the study as well as select structural ROIs (including hippocampus, entorhinal cortex) were preregistered in 2020 on https://aspredicted.org, but the methods used in the current paper were not pre-registered, as prolonged corona restrictions in part led to a much prolonged sample recruitment period, a sample of half the desired and registered size, and somewhat altered methods overall. Given this, along with methodological development in image processing, it was decided to use multi-feature cortical brainprints for the present study. We did not have hypotheses regarding brainprints in the initial preregistration, as we started using this methodology only later.

#### Cybercycle training program

The Cybercycle training program consisted of two variants designed to engage participants in navigating a virtual city called *Plasti City*: A Virtual Reality (VR) version conducted in our VR lab and an online/laptop-based version for playing remotely. These variants share similar tasks with slight differences in control schemes, and participants were asked to engage in both, ideally coming to the VR lab 2-3 times a week, and practicing the task at home 2-3 additional days a week. During the initial VR lab sessions, participants focused on adapting to the immersive VR environment. These sessions were deliberately kept brief at first and began with simple cycling along a virtual road, without any additional tasks (see below for details). Participants were encouraged to practice frequently but were advised to keep immersive VR sessions shorter (e.g., 20 minutes and no more than an hour) than the online or laptop-based sessions to reduce the risk of cyber-sickness (i.e., motion sickness). A series of tasks were presented to the participants, and unless otherwise noted, tasks were the same both in the VR lab and online/at laptops. Reporting on the results of these additional tasks is beyond the scope of the present paper, but for a full depiction of the intervention, they are also described below.

##### Description of the virtual city Plasti City

The virtual city used in the training program, Plasti City, is not based on any existing city and was manually built within Unreal Engine. The city is made up of several larger areas that are visually distinct, including a modern downtown in the south, a less dense suburb in the east, and a futuristic metropolis in the north. The total size of the city is roughly 5km² (Best estimate: 4.96km²). The north and south side of the city are split by a canal district. A coast follows the southern and western edges of the city.

##### Equipment

For immersive VR sessions, participants were seated on a standard mountain bike which was affixed to a FLUID^2 Indoor Bike Trainer. The FLUID^2 Trainer is a progressive resistance roller that allows for a realistic biking experience. For safety and stability during the experiment, participants were also equipped with a Woodway chest-shoulder harness. This harness, typically utilized for heart rate monitoring and safety during treadmill exercises, was modified for our purposes. It was securely attached to an overhead anchor point, allowing participants to maintain balance on the stationary bike without risk of falling. All virtual navigation tasks were conducted using two types of head-mounted display (HMD) systems: the HTC Vive Pro and the HTC Vive Cosmos Elite. The HTC Vive Pro HMD offers a high-resolution display (2880 x 1600 pixels) with a 110-degree field of view and a refresh rate of 90 Hz, whereas the HTC Vive Cosmos Elite features a combined pixel resolution of 2880 x 1700. Both HMDs provide a fully immersive 3D experience. Positional tracking was achieved through the SteamVR Tracking system, which ensures precision within the virtual space by using two base stations positioned in the testing area. A Vive controller attached to the bike’s handlebar and a Vive Tracker strapped inside the rear wheel of the bike allowed for tracking steering and wheel rotation respectively. All equipment was interfaced with a computer system equipped with an NVIDIA GeForce GTX 1080 Ti graphics card to render the virtual environment in real-time without perceptible latency. The virtual environment for the navigation tasks was developed using Unreal Engine 4.22.

#### Tasks in Plasti City

##### Main navigation task

In this task, participants are presented with a 3D model of a unique city building or feature. The model appears within a translucent globe, floating above the handlebar. It slowly rotates to allow participants to view it from all angles. The model remains visible until the participant reaches and successfully identifies the displayed target. To confirm target identification, participants must approach the target closely and press a button on the handlebar controller. The required distance is similar for all targets but may vary slightly. We have manually placed the “target area” to encompass the surrounding roads effectively (mean = 4.5 m, range: 3.5–6.9 m), see Figure S7. When a participant correctly identifies a target, their score is displayed on the screen. The scoring per trial, which is used for “leveling-up” in the game, represents the participant’s travel distance from the edge of the previous target’s area (or starting location for the first target) to the edge of the current target’s area, divided by the optimal distance between these points. The optimal distance is calculated using Unreal Engine’s native Navigation Mesh System. Scores of 1.3 or less (less than 30% over optimal distance) mark a successful trial, displayed in green with a consonant sound. Unsuccessful trials show the text in red with a dissonant sound. After identifying a target, the target model above the handlebar is replaced with the next target model.

##### Road Navigation

Participants can only navigate the city roads. If a participant veers off the road, the bike is instantly teleported back to its position and orientation from four meters ago. The bike’s speed before teleportation is maintained to minimize discomfort caused by simulated acceleration in VR.

##### Retrospective memory task

Starting in the fifth week of training for each group, the retrospective memory task is enabled. Participants receive a text description of the task, followed by a sequence of three navigation targets displayed alongside images of objects, for 5 seconds each. Participants must remember which targets were among the three and which specific images were shown. When they encounter one of the three targets during regular navigation, they must press a separate button from the main task. Incorrect button presses result in an error message, but there is no opportunity for correction. Mistakes still count as hits in the main navigation task. Correct button presses lead to a 3x3 grid of possible objects, one of which matches the initial image. In VR, participants select the target by turning their head toward the object, while the laptop version involves clicking with a mouse. Participants receive immediate feedback on the correctness of their answer. Six categories of images are used: Tools, fruit, hats, animals, furniture, and vehicles, each containing six subcategories, resulting in 216 possible target pictures. Picture selection depends on task difficulty, with eight levels: Difficulty Levels: 1. Six categories, 2. Three categories, 3. Two categories, 4. One category, 5. Three subcategories, 6. Two subcategories, 7. One subcategory, 8. One subcategory (grayscale). For example, at difficulty level 1, six pictures are selected from different categories. Level 2 involves two pictures from each of three categories. Level 4 uses pictures from the same category but different subcategories. Level 7 selects pictures from the same subcategory, while level 8 maintains the subcategory but presents images in grayscale. Task difficulty increases as participants perform well, ensuring challenges adapt to their progress. If participants ever had a 50% or higher accuracy on the previous six retrospective memory task presentations (counted across sessions), the difficulty level would increase by one for the next session. Starting difficulty was changed to level 4 based on preliminary data, indicating the task was too easy.

##### Passive Prospective Memory Task

Starting in the seventh week of training, the passive prospective memory task is enabled. In this task, many stationary 3D cars are introduced into the virtual city environment. The layout of these cars is unique for each session, generated using a randomly seeded procedural algorithm. The algorithm places the cars at semi-regular intervals on both sides of the road. We utilize different visual models for these cars, ensuring they are visually distinct from one another. After participants begin the experiment and receive instructions, they are shown one randomly selected car model that they must watch out for during regular navigation. This car model is displayed, slowly rotating in front of them for 10 seconds. Participants are instructed that if they encounter one of these target cars while navigating the city, they should press the designated target button. When pressing the target button while next to a car the participant gets shown a text prompt indicating whether they selected the correct car or not. If the participant passes by a target car without pressing a button, they receive a text prompt telling them they missed it.

##### Active Prospective Memory Task (Time-Based)

In the ninth week of training, the active prospective memory task becomes active. This task is exclusively available during virtual reality (VR) sessions and is not included in the at-home laptop-based sessions. In this task, participants are instructed to press a specific button on their VR controller when a certain amount of time has passed during a training session. The target time is randomized, ranging from a whole number between 3 and 9 minutes. This target time is displayed on a virtual screen within the VR environment each time a session begins. The current time in minutes and seconds is constantly displayed on the handlebar of the virtual bike. Starting in the tenth week, there is a modification to this task. The time display is no longer constantly visible but requires participants to press a separate button on the controller for it to appear. Participants are instructed to minimize the number of times they press this button, as each press results in a score reduction equivalent to a ten-second time penalty.

#### Difficulty Levels and “Leveling Up”

Both the main navigation task and the retrospective memory tasks were designed with adaptive difficulty. This means that the tasks become more challenging as participants perform well. In the main navigation task, increased difficulty entails expanding the navigable area and increasing the number of potential targets within the city. This is achieved by removing virtual barriers that initially block off certain streets. The city is divided into 13 areas, with the first unlocked at the start. Additional areas are unlocked based on participant performance. To trigger an increase in difficulty, a participant must reach three targets consecutively with a deviance from the path below the success criteria, as described below. Only targets from the latest unlocked area of the city are considered in this evaluation, ensuring that participants are sufficiently familiar with the current city area before more areas become accessible. To ensure that participants encounter new targets and have the chance to “level up” in any given session, at least three out of the first ten targets are always from the latest unlocked city level. In the retrospective memory task, difficulty (or “level”) is increased if participants correctly select three out of the latest five items across multiple sessions.

#### Behavioral data from Plasti City

Behavioral data from the tasks were collected online and directly from a set of identical laptops that participants borrowed and kept at home during the training period. Technical errors lead to a small portion of data known to be lost. Data from the task was saved in two ways: a long “continous log” of participants’ movement and input during the task and a shorted “event log”, only logging specific events such as button presses. There was a minor number of sessions for which the continous logs were saved, but not event logs enabling reconstruction of navigation and target count from the continous logs, and a few instances were observed where files lacked between levels, but the problem source could not be recreated. Participants in the project were initially able to log in and play online for their homework. However, due to experiences of some technical difficulties with online data collection for the homework tasks, it was after an initial period in the project decided that all participants should get identical laptops at which all data were automatically saved for collection post-training period. For the entire project period, 1189 online, 2304 laptop, and 1734 VR training sessions were recorded. Participants did themselves report on level reached in homework for correct placement of level each time they came to train in the VR lab. Homework data was retrieved from the laptops post-training. The levels saved from homework and the levels saved as trained in the VR-lab corresponded well (correlation of r =.94), and as a modest amount of data were lost due to technical difficulties, the highest level recorded played, whether that be at home or in the VR-lab, was used for analyses of “level reached”. However, for one set of DZ twins, there was no homework training data. These twins reported levels 4 and 3, respectively, and hence navigated accordingly at levels in VR. The pair was included as trainers in analyses, since they were recorded to have trained equal to the mean of the sample in immersive VR (4.27 and 5.39 hours of goal-directed navigation, targets counts: 242 and 226, mean relative deviance from shortest paths: 0.21 and 0.45, respectively). For all other training participants, both homework and immersive VR-data were retrieved and used in analyses of training times and level reached.

The measure of deviance from shortest path between two consecutive targets is estimated after transforming the city map into a skeletonized image, where all the streets are reduced to a 1-pixel-width representation. The shortest path is then determined using the Minimum Cost Path (MCP) approach, with the distance measured in pixels. The current participant’s path is also transformed into this same space, matching every position in the city to the closest skeletonized pixel. The deviance from shortest path is calculated by the difference between the participant’s path and the shortest path. The relative deviance from shortest path refers to the ratio between the deviance and the length of the shortest path. Both the skeleton transformation and the MCP algorithm are implemented using Python’s scikit-image package (https://doi.org/10.7717/peerj.453).

### Precision and Sensitivity of brainprint similarity

We leveraged data from N = 30 twin pairs (8 DZ), each with two passive scans (i.e. scans with no training in between), to assess the reliability of brainprint similarity. Intraclass correlation analysis indicated excellent reliability, with ICC(2,1) = 0.947 (CI: 0.891 – 0.974) (Figure S9A). The reliability was comparable when the analyses was restricted to MZ twins (N = 22 pairs) (ICC(2,1) = 0.929 (CI: 0.845 – 0.969)) (psych r-package). The within subject-measurement error – defined as the random variability in brainprint similarity across repeated scans – was also notably low (Sw= 0.0028) (Figure S9B). Given that brainprint similarity is a composite measure, these values reflect a remarkable level of precision. Additionally, we conducted a sensitivity analysis to determine the range of effect sizes that could be reliably detected given our sample size, corresponding statistical power, and a specified significance level. Sensitivity analysis, a form of power analysis particularly useful after data collection, allows for the estimation of minimum detectable effect sizes under realistic model assumptions. We generated sensitivity curves for two key analyses: (1) whether brainprint similarity at baseline is predicted by birth weight discordance × zygosity (representing early environmental effects; see Table S1: Model 1), and (2) whether brainprint similarity is influenced by the interaction between zygosity and training status (representing late environmental effects; see Table S1: Model 5). Specifically, Sensitivity curves were created using the simr r-package (n = 1000 simulations per model), with effect sizes increasing stepwise across simulations. Model structures and covariates were based on the empirical data, and we used linear models (lm [formula 1] and lmer [formula 5]) instead of generalized additive models (GAMs) for computational simplicity. In both cases, the interaction term was the effect of interest. See Sensitivity curves in Figure S9C-D. The sensitivity analysis showed that for both tests, the results have enough power, i.e, ≥ .8, to detect effects of medium size (η²□ ≈ .08 and .05 for the early and later environmental tests), which correspond roughly to raw effects on brainprint similarity of ≈0.0175 and 0.0023.

**Fig. S1.**
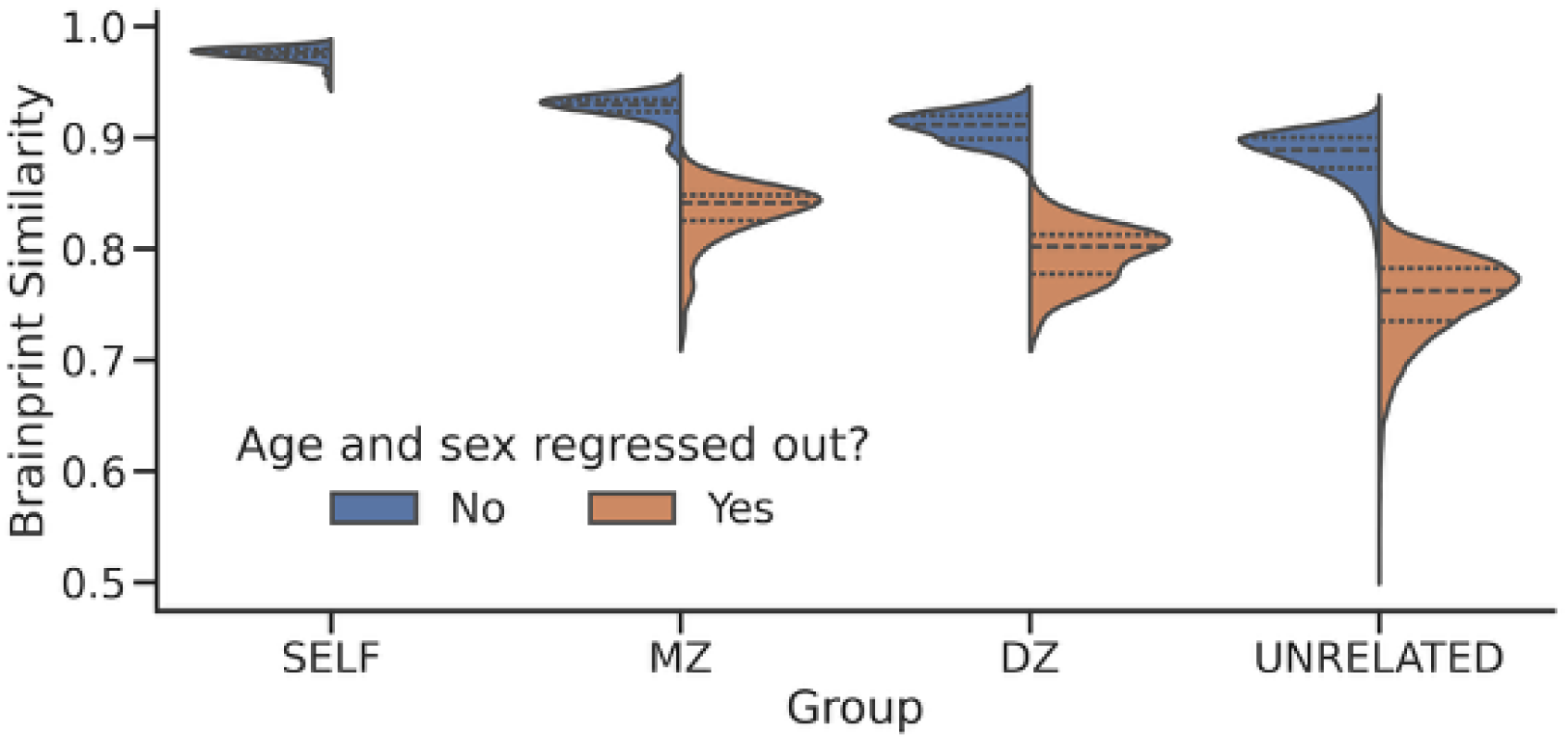
Distribution of brainprint similarity scores, both with and without regressing out age and sex from brain features going into the brainprints. The self-similarity distribution is here calculated from a subgroup of **127** individuals with two MRI sessions before training (passive non-trainers or “start_rest”-group), separated by about 10 weeks.

**Fig. S2.**
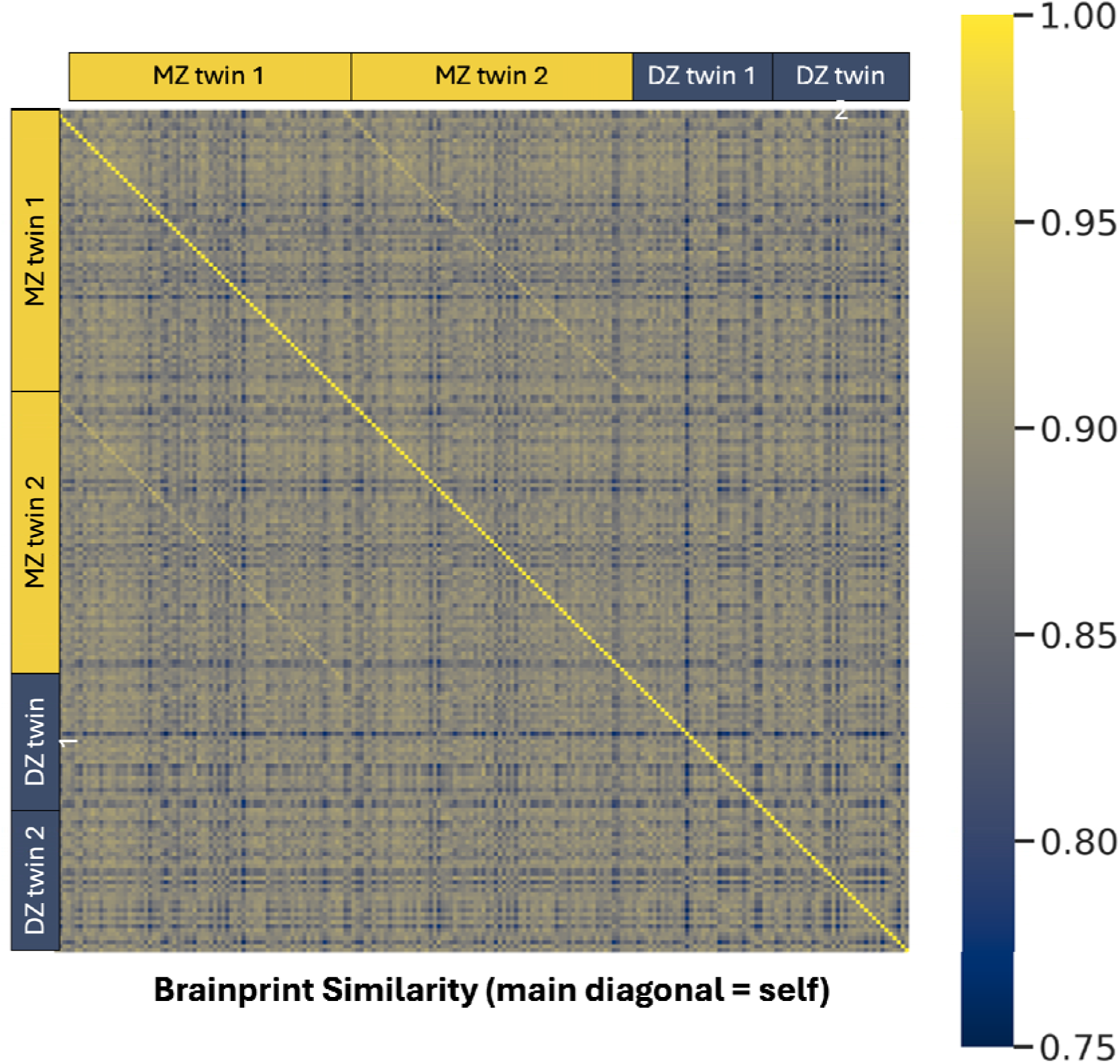
Raw brainprint similiarity at baseline, based on cortical area (CA), thickness (CT), volume (CV) and mean curvature (MC). Neither age, nor sex is controlled for here (compare to Figure 1A, showing brainprint similarity based on residuals when age and sex are controlled for). N = 210, 105 twin pairs (71 MZ, 34 DZ).

**Fig. S3.**
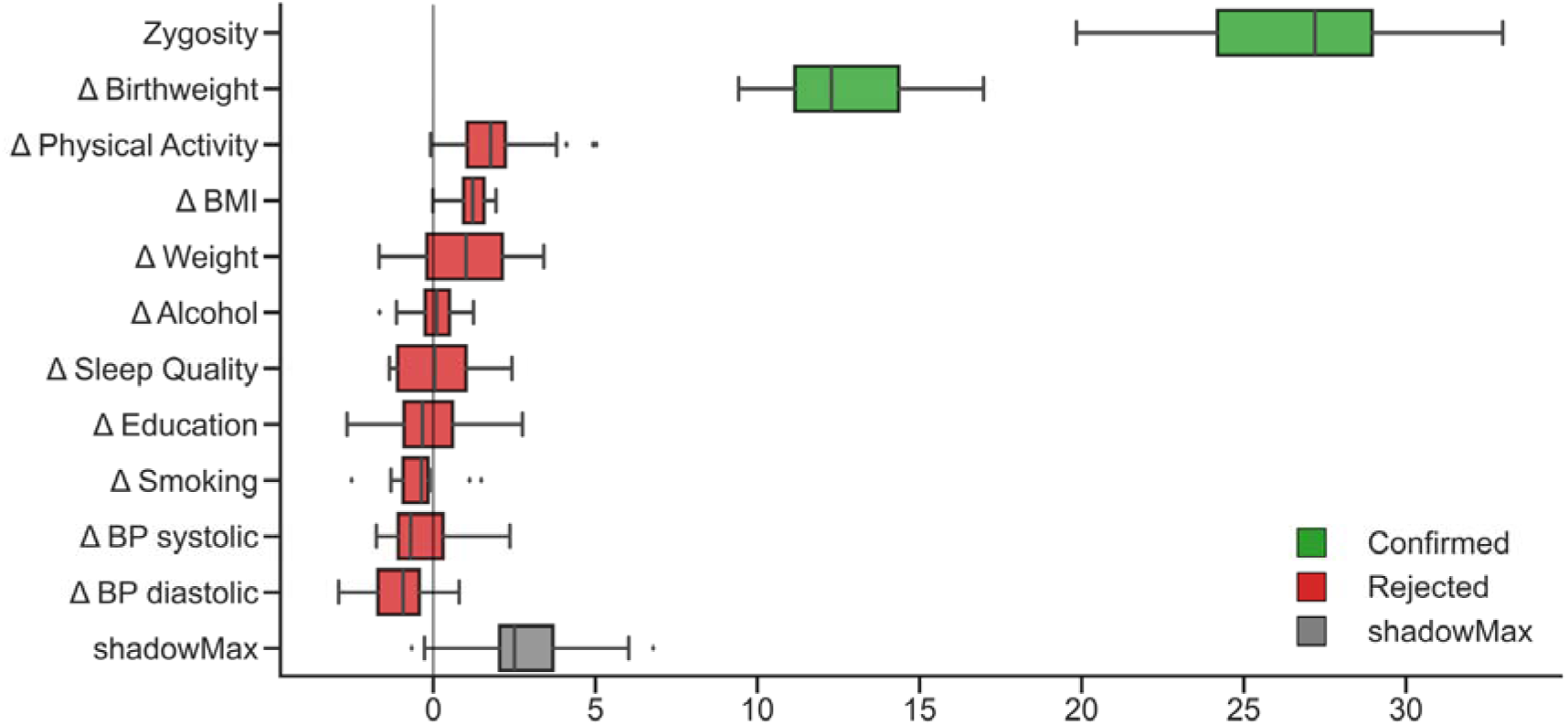
Feature selection using Random Forest (Boruta) on full sample of MZ and DZ twins. Zygosity comes out as the most important feature predicting brainprint similarity within twin pairs, followed by birthweight discordance. No other environmental discordance measures reached significance.

**Fig. S4.**
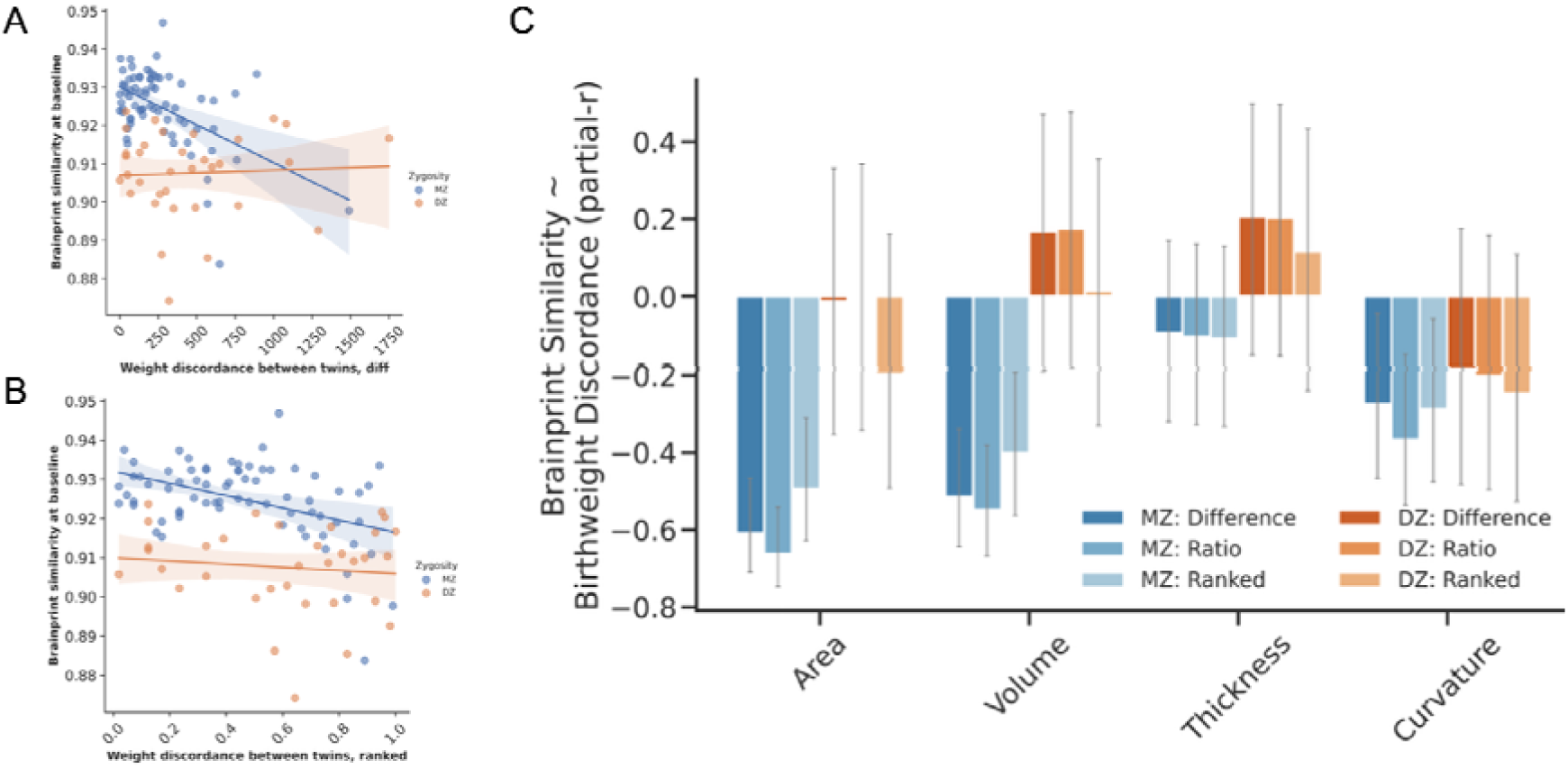
Birthweight discordance ranked. A) Same plot as main Figure 2B but with birthweight discordance operationalized as difference B) Simple ranking of weight discordance differences to achieve robust statistics. GAM Model 1 on ranked data produced similar conclusions as for the non-ranked operationalizations: brainprint similarity at baseline varied as a function of zygosity (t = -5.28, p < .0001) and weight discordance (t = -3.91, p = .0002). Weight discordance was significantly correlated with brainprint similarity in MZ (r = - .43, p = .0002), but not in DZ twins (r = -.11, p = .54). The interaction of zygosity and weight discordance was however not significant for ranked data (t = 1.706, p = .09). C) Same plot as main figure 2C, with data from other BW discordance operationalizations added. Cortical area and volume effects dominated also using ranked data (both p <= .001; curvature p = .018), and only in MZ twins (all DZ p > .10): increased MZ birthweight discordance (ranked) was associated with lower brainprint similarity in adulthood. Errorbars indicate 95% CI.

**Fig S5.**
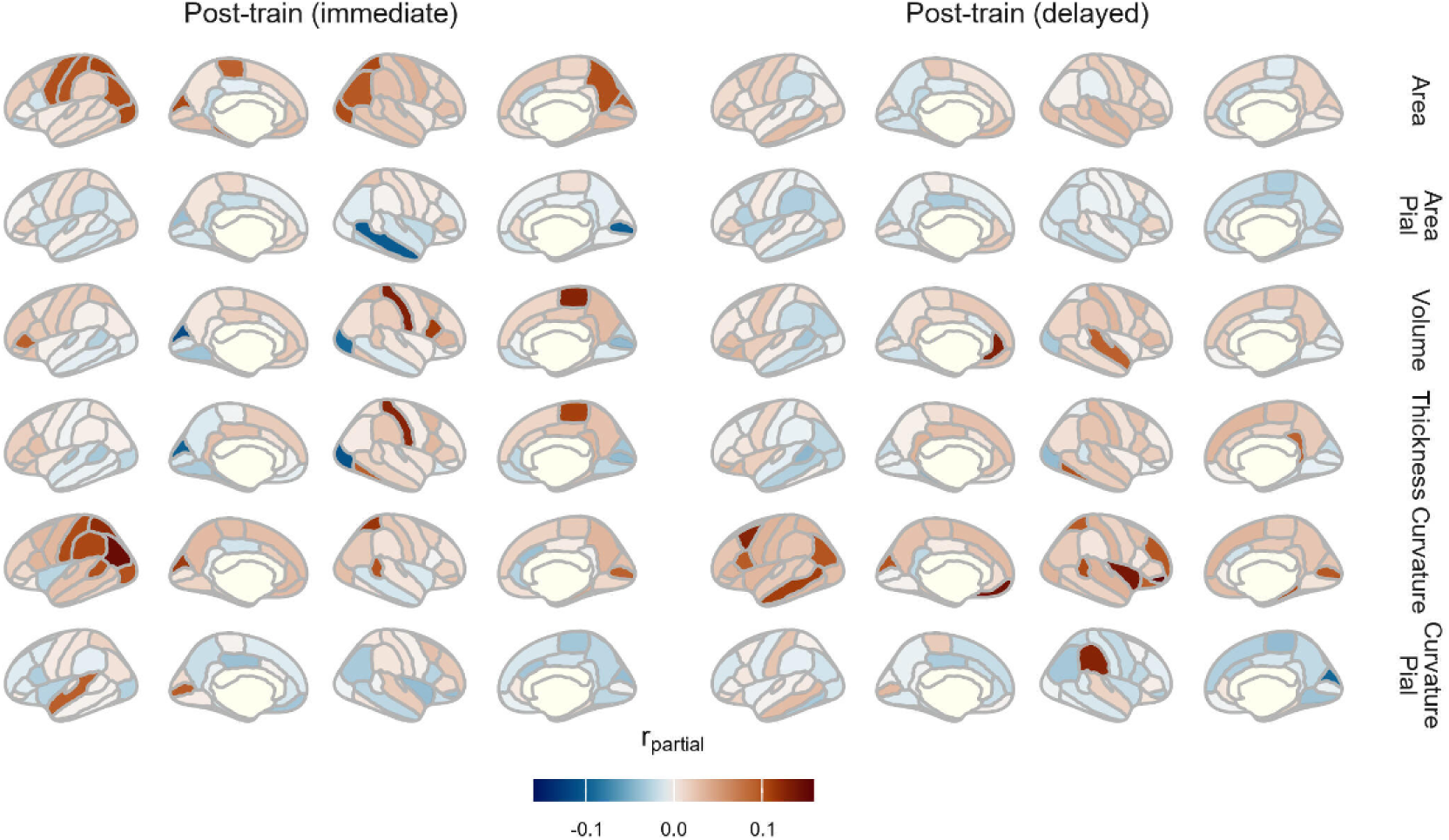
Effect of training status on regional cortical structure. The maps show partial correlation coefficients (r-partial) associated with training status (immediate and delayed post-train periods) and zygosity, from the GAMM model (Table S1, Model 6-7). Positive values indicate increase in regional cortical structure, i.e., relative increase in regional cortical area and curvature. Colours represent the partial correlation coefficient (r-partial) from the GAMM model, indicating the strength and direction of the association. Opaque regions are significant p < 0.05 (uncorrected). Note that no interpretation of these results is done at a regional level. The six rows correspond to different feature categories (Area, Area measured at the pial surface, Volume, Thickness, Curvature and Curvature measured at the pial surface). Brain views show left and right hemispheres, lateral and medial surfaces.

**Fig. S6.**
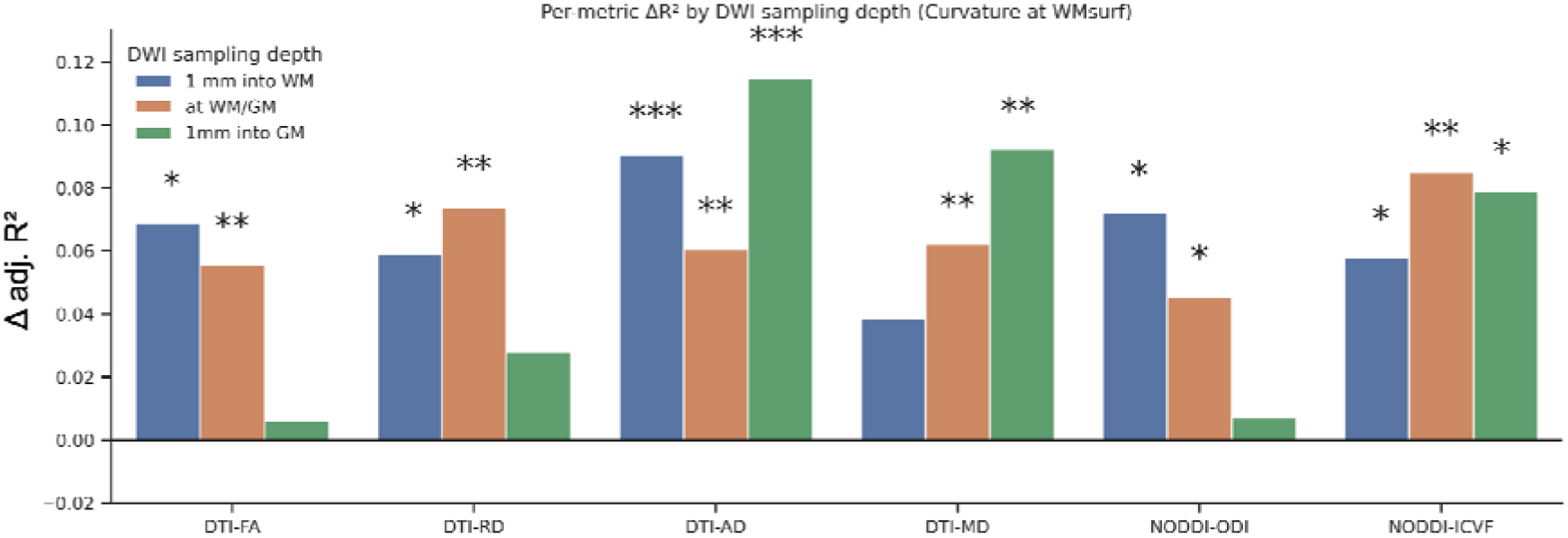
Per-metric improvement in predicting WM-surface curvature similarity when adding diffusion brainprints. For each diffusion metric a global twin-similarity predictor was computed at three depths (1 mm into WM, at WM/GM, 1 mm into GM) and added with its interaction with training to the model (Table S2: Model 8). Bars show change in adjusted R^2^ versus the baseline model without diffusion (Table S2: Model 5). Asterisks mark likelihood ratio test p-values after FDR correction across depths within each metric, evaluated at alpha = 0.05/6 due to six metrics (* p <.05, ** p < .01, *** p < .001).

**Fig. S7.**
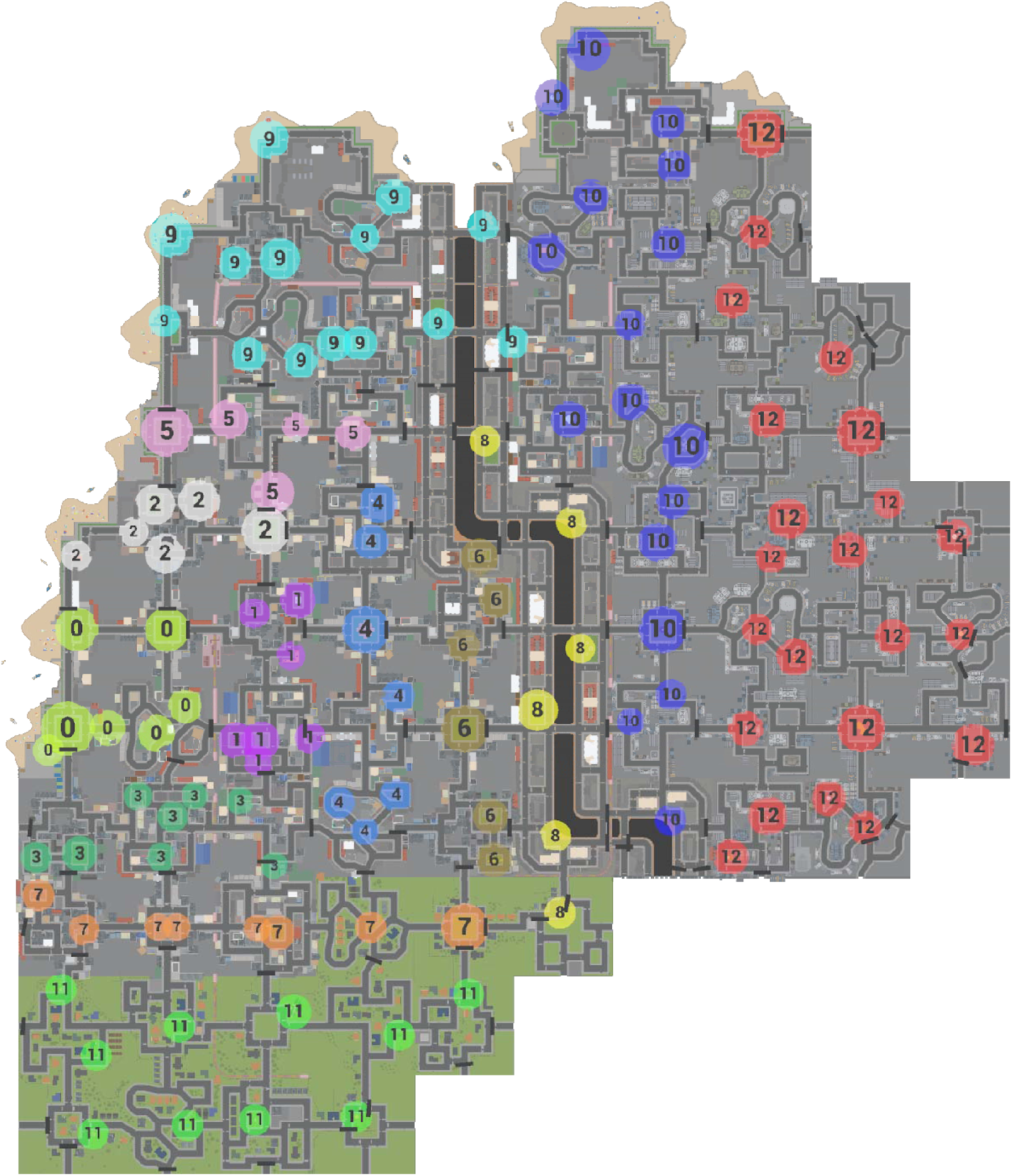
Distribution of targets, color coded per level, in Plasticity.

**Fig. S8.**
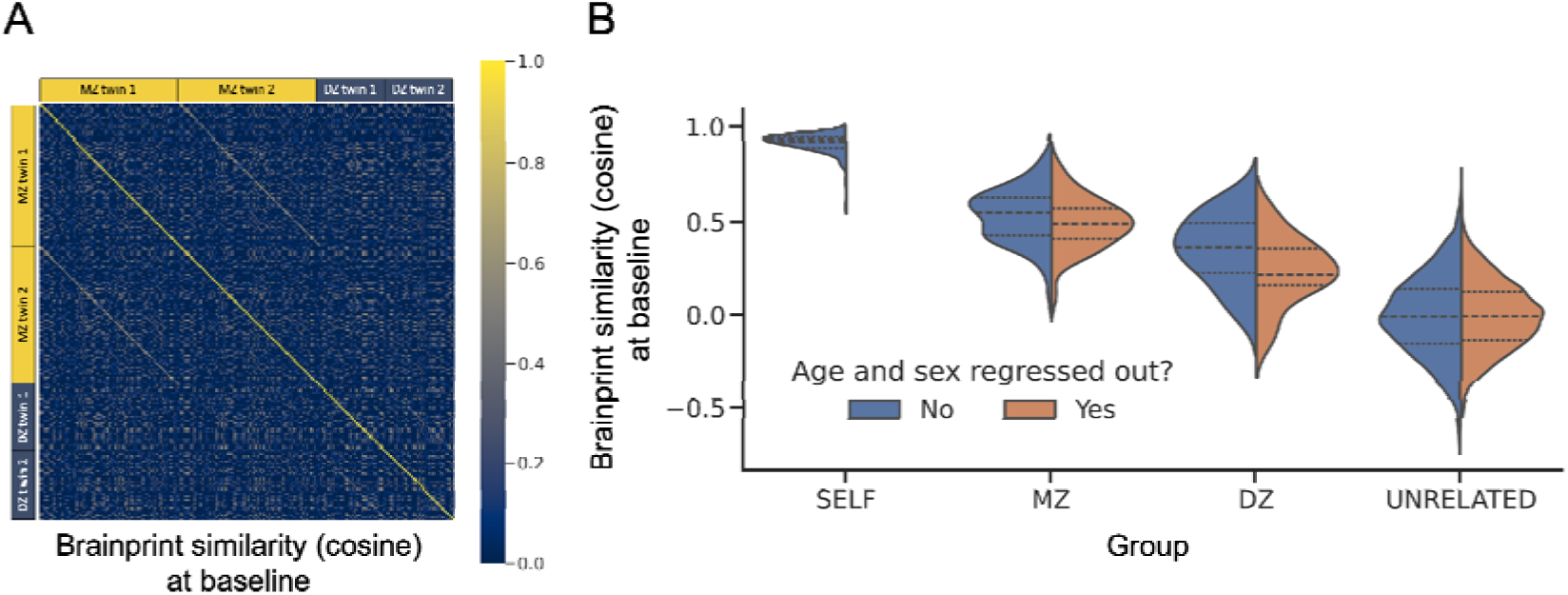
Cosine similarity as metric. A) Raw brainprint similarity calculated as for Fig. S2 but with cosine similarity as alternative metric. B) See caption to Fig. S1.

**Fig. S9.**
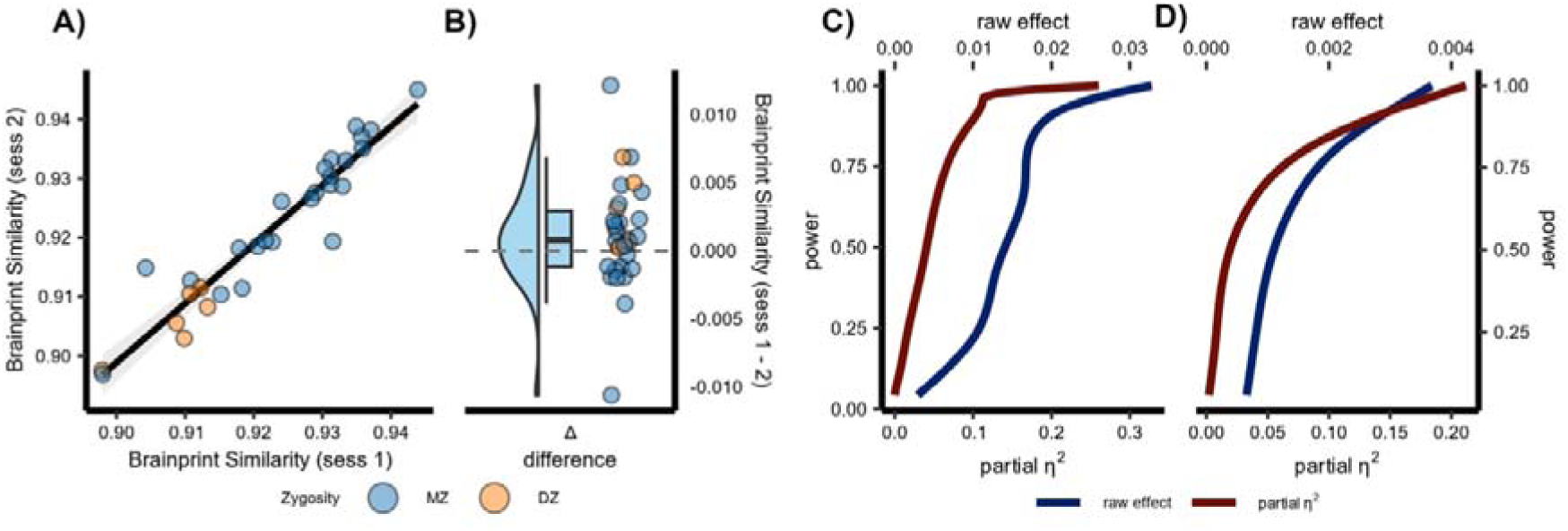
Precision and Sensitivity of brainprint similarity. Psychometric properties of brainprint similarity. A) Correlation of brainprint similarity between two different passive sessions (session 1 & session 2) for 30 twins (22 MZ). Colors represent zygosity. The line represents the correlation between session 1 and session 2 and the ribbon represents its standard error. B) Within-subject differences in brainprint similarity; i.e., random variability in brainprint similarity across repeated scans. Sensitivity analysis for the main C) early (BW discordance × zygosity) and D) late (training status × zygosity) environmental effects on brainprint similarity. The plots illustrate the relationship between effect size (raw, standardized) and power in our dataset.

## References

1. Grasby, K.L., et al. The genetic architecture of the human cerebral cortex. Science 367 (2020).

2. Rakic, P. Specification of cerebral cortical areas. Science 241, 170–176 (1988).

3. Desikan, R.S. & Barkovich, A.J. Malformations of cortical development. Ann Neurol 80, 797–810 (2016).

4. Eyler, L.T., et al. A comparison of heritability maps of cortical surface area and thickness and the influence of adjustment for whole brain measures: a magnetic resonance imaging twin study. Twin Res Hum Genet 15, 304–314 (2012).

5. Alexander-Bloch, A.F., et al. Imaging local genetic influences on cortical folding. Proceedings of the National Academy of Sciences of the United States of America 117, 7430–7436 (2020).

6. Snyder, W.E., et al. A bimodal taxonomy of adult human brain sulcal morphology related to timing of fetal sulcation and trans-sulcal gene expression gradients. Neuron (2024).

7. Storsve, A.B., et al. Differential longitudinal changes in cortical thickness, surface area and volume across the adult life span: regions of accelerating and decelerating change. The Journal of neuroscience : the official journal of the Society for Neuroscience 34, 8488–8498 (2014).

8. Draganski, B., et al. Neuroplasticity: changes in grey matter induced by training. Nature 427, 311–312 (2004).

9. Engvig, A., et al. Effects of memory training on cortical thickness in the elderly. Neuroimage 52, 1667–1676 (2010).

10. Engvig, A., et al. Effects of cognitive training on gray matter volumes in memory clinic patients with subjective memory impairment. Journal of Alzheimer’s disease : JAD 41, 779–791 (2014).

11. Wenger, E., et al. Cortical thickness changes following spatial navigation training in adulthood and aging. Neuroimage 59, 3389–3397 (2012).

12. Wenger, E., Brozzoli, C., Lindenberger, U. & Lovden, M. Expansion and Renormalization of Human Brain Structure During Skill Acquisition. Trends in cognitive sciences 21, 930–939 (2017).

13. Wenger, E., et al. Repeated Structural Imaging Reveals Nonlinear Progression of Experience-Dependent Volume Changes in Human Motor Cortex. Cereb Cortex 27, 2911–2925 (2017).

14. Olivo, G., et al. Estimated gray matter volume rapidly changes after a short motor task. Cereb Cortex 32, 4356–4369 (2022).

15. Federmann, L.M., et al. Associations between antagonistic SNPs for neuropsychiatric disorders and human brain structure. Transl Psychiatry 14, 406 (2024).

16. Radonjic, N.V., et al. Structural brain imaging studies offer clues about the effects of the shared genetic etiology among neuropsychiatric disorders. Mol Psychiatry 26, 2101–2110 (2021).

17. Thompson, P.M., et al. ENIGMA and global neuroscience: A decade of large-scale studies of the brain in health and disease across more than 40 countries. Transl Psychiatry 10, 100 (2020).

18. WHO. *Risk reduction of cognitive decline and dementia: WHO guidelines* (World Health Organization, Geneva, 2019).

19. Livingston, G., et al. Dementia prevention, intervention, and care: 2024 report of the Lancet standing Commission. Lancet 404, 572–628 (2024).

20. Wheater, E., et al. Birth weight is associated with brain tissue volumes seven decades later but not with MRI markers of brain ageing. Neuroimage Clin 31, 102776 (2021).

21. Walhovd, K.B., et al. Neurodevelopmental origins of lifespan changes in brain and cognition. Proceedings of the National Academy of Sciences of the United States of America 113, 9357–9362 (2016).

22. Walhovd, K.B., et al. Long-term influence of normal variation in neonatal characteristics on human brain development. Proceedings of the National Academy of Sciences of the United States of America 109, 20089–20094 (2012).

23. Walhovd, K.B., et al. Fetal influence on the human brain through the lifespan. Elife 12 (2024).

24. Brouwer, R.M., et al. Genetic influences on individual differences in longitudinal changes in global and subcortical brain volumes: Results of the ENIGMA plasticity working group. Hum Brain Mapp 38, 4444–4458 (2017).

25. Walhovd, K.B., Lovden, M. & Fjell, A.M. Timing of lifespan influences on brain and cognition. Trends in cognitive sciences 27, 901–915 (2023).

26. Assary, E., et al. Genetics of monozygotic twins reveals the impact of environmental sensitivity on psychiatric and neurodevelopmental phenotypes. Nat Hum Behav 9, 1683–1696 (2025).

27. Fox, P.W., Hershberger, S.L. & Bouchard, T.J., Jr. Genetic and environmental contributions to the acquisition of a motor skill. Nature 384, 356–358 (1996).

28. Elmer, S., Hanggi, J., Meyer, M. & Jancke, L. Increased cortical surface area of the left planum temporale in musicians facilitates the categorization of phonetic and temporal speech sounds. Cortex 49, 2812–2821 (2013).

29. Fjell, A.M., et al. Continuity and Discontinuity in Human Cortical Development and Change From Embryonic Stages to Old Age. Cereb Cortex 29, 3879–3890 (2019).

30. Hogstrom, L.J., Westlye, L.T., Walhovd, K.B. & Fjell, A.M. The structure of the cerebral cortex across adult life: age-related patterns of surface area, thickness, and gyrification. Cereb Cortex 23, 2521–2530 (2013).

31. Huttenlocher, P.R. Synaptic density in human frontal cortex - developmental changes and effects of aging. Brain research 163, 195–205 (1979).

32. Manuello, J., et al. The effects of genetic and modifiable risk factors on brain regions vulnerable to ageing and disease. Nat Commun 15, 2576 (2024).

33. Walhovd, K.B. & Lövdén, M. A lifespan perspective on human neurocognitive plasticity. in The Cognitive Neurosciences (ed. Gazzaniga, Mangun & Poeppel) (MIT Press, Cambridge, MA, 2019).

34. Walhovd, K.B., Storsve, A.B., Westlye, L.T., Drevon, C.A. & Fjell, A.M. Blood markers of fatty acids and vitamin D, cardiovascular measures, body mass index, and physical activity relate to longitudinal cortical thinning in normal aging. Neurobiol Aging 35, 1055–1064 (2014).

35. Casey, K.F., et al. Birth weight discordance, DNA methylation, and cortical morphology of adolescent monozygotic twins. Hum Brain Mapp 38, 2037–2050 (2017).

36. Raznahan, A., Greenstein, D., Lee, N.R., Clasen, L.S. & Giedd, J.N. Prenatal growth in humans and postnatal brain maturation into late adolescence. Proceedings of the National Academy of Sciences of the United States of America 109, 11366–11371 (2012).

37. Mediavilla, T., et al. Learning-related contraction of gray matter in rodent sensorimotor cortex is associated with adaptive myelination. Elife 11 (2022).

38. Sebenius, I., et al. Structural MRI of brain similarity networks. Nature reviews. Neuroscience (2024).

39. Valizadeh, S.A., Liem, F., Merillat, S., Hanggi, J. & Jancke, L. Identification of individual subjects on the basis of their brain anatomical features. Sci Rep 8, 5611 (2018).

40. Wachinger, C., et al. BrainPrint: a discriminative characterization of brain morphology. Neuroimage 109, 232–248 (2015).

41. Desikan, R.S., et al. An automated labeling system for subdividing the human cerebral cortex on MRI scans into gyral based regions of interest. Neuroimage 31, 968–980 (2006).

42. Pienaar, R., Fischl, B., Caviness, V., Makris, N. & Grant, P.E. A Methodology for Analyzing Curvature in the Developing Brain from Preterm to Adult. Int J Imaging Syst Technol 18, 42–68 (2008).

43. Buysse, D.J., Reynolds, C.F., 3rd, Monk, T.H., Berman, S.R. & Kupfer, D.J. The Pittsburgh Sleep Quality Index: a new instrument for psychiatric practice and research. Psychiatry Res 28, 193–213 (1989).

44. Craig, C.L., et al. International physical activity questionnaire: 12-country reliability and validity. Med Sci Sports Exerc 35, 1381–1395 (2003).

45. Coughlan, G., et al. Toward personalized cognitive diagnostics of at-genetic-risk Alzheimer’s disease. Proceedings of the National Academy of Sciences of the United States of America 116, 9285–9292 (2019).

46. Spiers, H.J., Coutrot, A. & Hornberger, M. Explaining World-Wide Variation in Navigation Ability from Millions of People: Citizen Science Project Sea Hero Quest. Top Cogn Sci 15, 120–138 (2023).

47. Lovden, M., et al. Spatial navigation training protects the hippocampus against age-related changes during early and late adulthood. Neurobiol Aging 33, 620 e629–620 e622 (2012).

48. Walhovd, K.B., et al. Within-session verbal learning slope is predictive of lifespan delayed recall, hippocampal volume, and memory training benefit, and is heritable. Sci Rep 10, 21158 (2020).

49. Kursa, M.B. & Rudnicki, W.R. Feature Selection with the Boruta Package. Journal of Statistical Software 26, 1–13 (2010).

50. Halevy, T., et al. Twin discordance: a study of volumetric fetal brain MRI and neurodevelopmental outcome. Eur Radiol 31, 6676–6685 (2021).

51. Mukherjee, N., et al. Discordance of prenatal and neonatal brain development in twins. Early Hum Dev 85, 171–175 (2009).

52. Walhovd, K.B., Fjell, A.M., Giedd, J., Dale, A.M. & Brown, T.T. Through Thick and Thin: a Need to Reconcile Contradictory Results on Trajectories in Human Cortical Development. Cereb Cortex 27, 1472–1481 (2017).

53. (!!! INVALID CITATION !!! 31).

54. Greenough, W.T., Black, J.E. & Wallace, C.S. Experience and brain development. Child development 58, 539–559 (1987).

55. Petanjek, Z., et al. Extraordinary neoteny of synaptic spines in the human prefrontal cortex. Proceedings of the National Academy of Sciences of the United States of America 108, 13281–13286 (2011).

56. Yeung, M.S., et al. Dynamics of oligodendrocyte generation and myelination in the human brain. Cell 159, 766–774 (2014).

57. Bhardwaj, R.D., et al. Neocortical neurogenesis in humans is restricted to development. Proceedings of the National Academy of Sciences of the United States of America 103, 12564–12568 (2006).

58. Paredes, M.F., Sorrells, S.F., Garcia-Verdugo, J.M. & Alvarez-Buylla, A. Brain size and limits to adult neurogenesis. J Comp Neurol 524, 646–664 (2016).

59. Rehn, A.E., et al. An animal model of chronic placental insufficiency: relevance to neurodevelopmental disorders including schizophrenia. Neuroscience 129, 381–391 (2004).

60. Scarr-Salapatek, S. Race, social class, and IQ. Science 174, 1285–1295 (1971).

61. Tucker-Drob, E.M. & Bates, T.C. Large Cross-National Differences in Gene x Socioeconomic Status Interaction on Intelligence. Psychol Sci 27, 138–149 (2016).

62. Rash, B.G., Arellano, J.I., Duque, A. & Rakic, P. Role of intracortical neuropil growth in the gyrification of the primate cerebral cortex. Proceedings of the National Academy of Sciences of the United States of America 120, e2210967120 (2023).

63. Natu, V.S., et al. Apparent thinning of human visual cortex during childhood is associated with myelination. Proceedings of the National Academy of Sciences of the United States of America 116, 20750–20759 (2019).

64. Herrera-Luis, E., Benke, K., Volk, H., Ladd-Acosta, C. & Wojcik, G.L. Gene-environment interactions in human health. Nat Rev Genet 25, 768–784 (2024).

65. Sidman, R.L. & Rakic, P. Neuronal migration, with special reference to developing human brain: a review. Brain research 62, 1–35 (1973).

66. Nilsen, T., Brandt, I. & Harris, J.R. The Norwegian Twin Registry. Twin Res Hum Genet 22, 647–650 (2019).

67. Walhovd, K.B., et al. Genetic risk for Alzheimer disease predicts hippocampal volume through the human lifespan. Neurol Genet 6, e506 (2020).

68. Chang, C.C., et al. Second-generation PLINK: rising to the challenge of larger and richer datasets. Gigascience 4, 7 (2015).

69. Nilsen, T.S., Brandt, I., Magnus, P. & Harris, J.R. The Norwegian Twin Registry. Twin Res Hum Genet 15, 775–780 (2012).

70. Nilsen, T.S., Kutschke, J., Brandt, I. & Harris, J.R. Validity of Self-Reported Birth Weight: Results from a Norwegian Twin Sample. Twin Res Hum Genet 20, 406–413 (2017).

71. Ji, J.L., et al. QuNex-An integrative platform for reproducible neuroimaging analytics. Front Neuroinform 17, 1104508 (2023).

72. Glasser, M.F., et al. The minimal preprocessing pipelines for the Human Connectome Project. Neuroimage 80, 105–124 (2013).

73. Zhang, H., Schneider, T., Wheeler-Kingshott, C.A. & Alexander, D.C. NODDI: practical in vivo neurite orientation dispersion and density imaging of the human brain. Neuroimage 61, 1000–1016 (2012).

74. Hernandez-Fernandez, M., et al. Using GPUs to accelerate computational diffusion MRI: From microstructure estimation to tractography and connectomes. Neuroimage 188, 598–615 (2019).

75. Fukutomi, H., et al. Neurite imaging reveals microstructural variations in human cerebral cortical gray matter. Neuroimage 182, 488–499 (2018).

76. Puonti, O., Iglesias, J.E. & Van Leemput, K. Fast, sequence adaptive parcellation of brain MR using parametric models. Med Image Comput Comput Assist Interv 16, 727–734 (2013).

77. Puonti, O., Iglesias, J.E. & Van Leemput, K. Fast and sequence-adaptive whole-brain segmentation using parametric Bayesian modeling. Neuroimage 143, 235–249 (2016).

78. Sebenius, I., et al. Robust estimation of cortical similarity networks from brain MRI. Nat Neurosci 26, 1461–1471 (2023).

79. Madan, C.R. Robust estimation of sulcal morphology. Brain Inform 6, 5 (2019).

80. Pedregosa, F., et al. Scikit-learn: Machine Learning in Python. Journal of Machine Learning Research 12, 2825–2830 (2011).

81. Nilearn contributors, et al. Zenodo. (2025).

